# *Medicago truncatula* ABCG40 is a cytokinin importer that negatively regulates lateral root density and nodule number

**DOI:** 10.1101/2022.11.10.516000

**Authors:** Tomasz Jamruszka, Joanna Banasiak, Aleksandra Pawela, Karolina Jarzyniak, Jian Xia, Wanda Biała-Leonhard, Lenka Plačková, Tashi Tsering, Francesca Romana Iacobini, Ondřej Novák, Markus Geisler, Michał Jasiński

## Abstract

Numerous studies suggest that cytokinin (CK) distribution plays a relevant role in shaping plant morphology in changing environments. Nonetheless, our knowledge about the involvement of short-distance CK translocation in root mineral nutrition remains scarce, and the specific role of CK transporters in root morphology has yet to be established. Therefore, the molecular identity of CK transporters should be determined to increase knowledge on root plasticity during soil fertility, as well as more frequently encountered plant nutrient deficiencies. In this work, we identified and characterized the *Medicago truncatula* full-size ATP-binding cassette (ABC) transporter of the G subfamily MtABCG40 as a plasma membrane CK importer. Its expression is root-specific and is induced by nitrogen deprivation and CKs. Our analyses indicate that MtABCG40 exerts a negative impact on lateral root density by decreasing lateral root initiation and enhancing primary root elongation. Moreover, we also observed that this transporter negatively influenced the nodule number. Our results suggest that MtABCG40 action affects CK signalling, which impacts the cellular response to auxin. In summary, we identified a novel ABCG-type CK transporter that regulates lateral root density and nodule number.

## Introduction

As the availability of nitrogen in soil fluctuates, plants can adapt their physiology and morphology accordingly to meet their needs, which is aided by the high plasticity of their roots. Changes in root physiology are controlled by the production and translocation of various signalling molecules. These molecules include phytohormones, such as cytokinins (CKs), that can exert local effects or trigger a systemic response. These effects alter root morphology together with nitrogen uptake from the soil (Jia and von Wiren, 2020). In reactions to varying nitrogen concentrations, plants can adjust the primary root length and lateral root number, which is often manifested by changes in lateral root density (Lopez-Bucio et al., 2003; Lima et al., 2010; Postma et al., 2014). An increase in the distance between adjacent lateral roots, called lateral root spacing, is triggered by different factors; one prominent factor is CKs, which suppress a positive effect of auxin on lateral root initiation (Laplaze et al., 2007). The spacing also results from the influence of CKs on the root apical meristem (RAM), as the size and root growth are decelerated (Dello Ioio et al., 2007; Dello Ioio et al., 2008). In legumes, a change in root morphology during nitrogen deficiency can also result from an interaction with symbiotic partners, namely, nitrogen-fixing soil bacteria collectively called rhizobia. The latter induce cortical cell divisions to form root nodules (Oldroyd et al., 2011). Notably, CKs in legumes control the initial steps of nodule formation by promoting cell proliferation in the cortex, generating nodule primordia (Gonzalez-Rizzo et al., 2006; Murray et al., 2007). However, CKs can also act as negative regulators of nodulation by inhibiting further infections in the epidermis (Miri et al., 2019) and systemically suppressing primordium formation (Sasaki et al., 2014). Of note, CK action in lateral root and nodule organogenesis is highly dependent on the nitrogen status of plants (Gu et al., 2018).

Aliphatic CKs are adenine derivatives with isoprenoid substitutions at the N^6^ position. Ribosides, their biologically inactive forms, exhibit more complex structures with an additional ribose moiety and are translocated along plant within vascular tissues (Kieber and Schaller, 2014; Osugi et al., 2017). The cleavage of the sugar component from CK riboside 5’- monophosphates by LONELY GUY (LOG) enzymes leads to the formation of their active forms, isopentenyladenine (iP) or *trans*-zeatin (*t*Z) (Kurakawa et al., 2007). To trigger specific outcomes, CKs must be translocated across biological membranes and perceived, either intracellularly or extracellularly (Romanov et al., 2018).

To date, several CK transporters with functions in roots have been described. Three of them, namely, MtABCG56, AtABCG14, and OsABCG18, belong to the G subfamily of the ATP-BINDING CASSETTE (ABC) family of transporters (Ko et al., 2014; Zhang et al., 2014; Zhao et al., 2019; Jarzyniak et al., 2021). ABCG transporters in general translocate molecules (as exporters and importers) across biological membranes using ATP as a source of energy. Their action has been assigned to different developmental processes, reactions to abiotic stresses, interactions with pathogens, and symbiotic associations (Lefevre and Boutry, 2018). These pathways also involve the translocation of other phytohormones, such as strigolactones (Kretzschmar et al., 2012; Banasiak et al., 2020) and abscisic acid (ABA) (Kang et al., 2010; Pawela et al., 2019). To date, the only CK transporter implicated in nodulation, MtABCG56, is localized to the plasma membrane (PM) and transports *t*Z as well as iP. MtABCG56 exports CK from rhizodermal and cortical cells after sensing symbiotic bacteria-derived Nod factors in the root susceptible zone (Jarzyniak et al., 2021). On the other hand, *ABCG14* from nonsymbiotic *A. thaliana* is expressed mainly in the pericycle and vasculature of the root along with CK biosynthesis genes, such as *ISOPENTENYLTRANSFERASE 3 (IPT3)* and *CYTOCHROME P450 (CYP) MONOOXYGENASES*. AtABCG14 is a PM *t*Z-type CK efflux pump that contributes to its long-distance translocation throughout the xylem and subsequent systemic impact on plant development (Ko et al., 2014; Zhang et al., 2014). Notably, a similar function was later demonstrated for its orthologue from *Oryza sativa*, OsABCG18 (Zhao et al., 2019).

Other CK transporters known to function in roots belong to the PURINE PERMEASE (PUP), AZA-GUANINE RESISTANCE (AZG) and EQUILIBRATIVE NUCLEOSIDE TRANSPORTER (ENT) families. AtPUP14 is a PM importer of bioactive CKs (*t*Z, iP and 6- benzylaminopurine 6-BAP) in Arabidopsis seedlings, specifically in their root tip meristematic cells and lateral root primordia. As PUP14 activity creates a sink for CKs inside the cell, the hormone can no longer be perceived in the apoplast by PM-bound receptors. Loss of PUP14 transport activity leads to root morphological defects (Zurcher et al., 2016). AtAZG2, which localizes to the PM and endoplasmic reticulum (ER) of root cells, transports iP, *t*Z, 6-BAP and kinetin. *AtAZG2* is expressed in tissues overlaying lateral root primordia and, as opposed to auxin, negatively influences lateral root emergence (Tessi et al., 2021). Finally, AtENT3 translocates CK nucleotides through the PMs of root vascular bundles and in the root tip, enabling the hormone to systematically move throughout the plant and affect its development (Traub et al., 2007; Cornelius et al., 2012; Korobova et al., 2021).

Here, we identified and characterized the *Medicago truncatula* full-size ABCG transporter MtABCG40 as a CK importer. *MtABCG40* expression is root-specific and is induced by nitrogen deprivation and CKs. Our analyses indicate that MtABCG40 exerts a negative impact on lateral root density through decreasing lateral root initiation and enhancing primary root elongation. Moreover, a lack of this transporter leads to an increased nodule number. Our results suggest that MtABCG40 action affects CK signalling, which impacts the cellular response to auxin.

## Materials and Methods

### Plant materials and growth conditions

*Medicago truncatula Tnt1* retrotransposon insertion mutant lines, namely NF21323 (*mtabcg40- 1*) and NF17891 (*mtabcg40-2*), were obtained from the Noble Research Institute. The presence of the respective insertions was confirmed using polymerase chain reaction (PCR) with *Tnt1*- and gene-specific primers (Table S1A). The level of *MtABCG40* expression in homozygous plants was verified using quantitative reverse transcription PCR (RT–qPCR) with gene-specific primers (Table S1A).

The seeds of *M. truncatula* Jemalong A17, R108, R108/DR5:GUS stable transgenic plants and *mtabcg40* plants were scarified with 96% sulfuric acid for 10 min, stratified on 0.8% agar plates for 3 days at 4°C and germinated overnight at 21°C. The seedlings were then grown in growth chambers under a 16 h light/8 h dark regime, at 22°C and 50%–60% relative humidity.

### Bacterial strains and growth conditions

Rhizobial strains, namely *Sinorhizobium meliloti* 1021 (wild-type strain), *S. meliloti* SL44 (1021 strain with deletion in the nod genes *ΔnodD1ABC*), *S. meliloti* E65 (A2101/pE65; 1021 strain containing pE65 plasmid that constitutively expresses the *nodD3* gene) and *S. meliloti* Rm1021/pXLGD4 (1021 strain containing plasmid that constitutively expresses the *lacZ* gene) were used for the inoculation of *M. truncatula*. Bacteria were grown in Bergensen’s modified medium (BMM) (Rolfe et al., 1980) containing appropriate antibiotics or 3 μM luteolin (prior to spot-inoculation assay).

### Plant treatments

For lateral root density measurement, seedlings after germination (NH_4_NO_3_ and 6-BAP gradients, *mtabcg40* mutants experiments), 1-week-old seedlings (IAA gradient experiment) or 1-week-old composite plants (*MtABCG40* and *MtLOG3* RNAi silencing experiments) grown initially on full-strength Fähraeus medium were then transferred directly onto modified Fähraeus (Barker et al., 2006) agar plates not supplemented with nitrogen (here referred to as 0 mM NH_4_NO_3_), with different NH_4_NO_3_ concentrations or full-strength Fähraeus agar plates with varying hormone contents, depending on the type of experiment. The plants were grown for one (IAA gradient experiment), two (6-BAP gradient experiment) or three (NH_4_NO_3_ gradient, *mtabcg40* mutants experiments) weeks, after which the root measurements were conducted. Lateral root density was calculated by dividing the number of first order lateral roots by the length of primary root. Analyses were performed on ≥31 (31-45) roots per condition in three independent experimental repetitions. If specified, the whole root samples were collected and immediately frozen for expression analyses. Roots from three plants were pooled for each condition in qRT-PCR analyses, in three biological replicates.

For iP and *t*Z treatments, 7-day-old *M. truncatula* seedlings grown on solid half-strength Murashige and Skoog medium (½ MS; M5524, Sigma-Aldrich) were transferred onto fresh ½ MS medium supplemented with 1 µM of an appropriate hormone, dissolved in NaOH or the equal volume of NaOH (mock). For expression analyses, whole roots were collected and immediately frozen after 1, 6, and 24 h of treatment. Three independent replicates for each time point were performed with three roots collected per sample.

For the expression analyses in RAM, root apical fragments of mutant (*mtabcg40-1*) and corresponding WT plants grown for 10 days on 0 or 1 mM NH_4_NO_3_ Fähraeus agar plates were collected. Three independent replicates were performed with ∼20 RAMs collected per sample.

For gravity-stimulated lateral root initiation, plants were grown vertically on 0.5 or 1.0 mM NH_4_NO_3_ Fähraeus agar plates for three days, turned 135° for 12 h to constrain a growth-mediated curvature of the root, and turned back afterward to the initial orientation. Root samples, comprising 10-15 newly created < 0.5 cm bents fragments each, were collected 12, 24, and 48 h after onset of lateral root initiation and immediately frozen for expression analyses. Three independent biological repeats were performed.

For *S. meliloti* inoculation assays, seedlings after germination or 3-week-old composite plants grown on full-strength Fähraeus medium were transferred directly onto 0 mM NH_4_NO_3_ Fähraeus agar plates (expression analyses and EdU staining experiments) or into pots (0.5 l) containing vermiculite/perlite/soil (3:2:1.5, v/v) and supplemented with 0 mM NH_4_NO_3_ Fähraeus medium twice a week (promoter activity analyses, assessment of nodule number experiments). For expression analyses with *S. meliloti* SL44 and E65 roots of 4-day-old Jemalong A17 plants were flood-inoculated with 200 µl of bacterial suspension (OD_600_ =0.01). Whole root samples were then collected and immediately frozen for expression analyses at the specified time points. For spot-inoculation assays, a 0.5 µl droplet of *S. meliloti* 2011 strain (OD_600_=0.02) was applied on the root susceptible zone (where the root hairs appear and still elongate) of each of 3-day-old seedlings. 8-12 root fragments (2-3 mm each) comprising the spot-inoculated area were collected 12, 24, and 48 h after an application of a droplet and immediately frozen for expression analyses. For assessment of nodule number and analysis of promoter activity, roots were inoculated with 50 ml of *S. meliloti* 1021 or *S. meliloti* Rm1021/pXLGD4 strain suspension (OD_600_ = 0.01) per pot, respectively. Nodule number was counted 21 days post-inoculation (dpi), for GUS staining the root material was collected from 5 to 21 dpi. Each *S. meliloti* inoculation assay was performed in two or three independent biological repeats.

### Genetic constructs and plant transformation

For the analysis of tissue-specific expression pattern, 2015 bp and 2073 bp fragments upstream of ATG start codon, corresponding to the promoter regions of *MtABCG40* and *MtLOG3*, respectively, were amplified with KOD Hot Start DNA Polymerase (Novagen) and cloned into pDONR/Zeo vector (Thermo Fisher) using Gateway Recombination Cloning Technology (Reece-Hoyes and Walhout, 2018). The obtained entry clone was subsequently recombined with pKGWFS7 destination vector, containing the GUS reporter gene sequence (Karimi et al., 2002). *MtABCG40* and *MtLOG3* promoters were also amplified and cloned through ligation-independent cloning into pPLV04_v2 (*MtABCG40:NLS-GFP*) and pPLV11_v2 (*MtLOG3:NLS-tdTomato*) vectors, respectively, both carrying reporter genes tagged with a nuclear localization signal (SV40) (De Rybel et al., 2011). Cloning of *MtRR4* promoter region (2118 bp) to pKGWFS7 vector was previously described by Jarzyniak et al. (2021). A list of primers can be found in Table S1B.

For subcellular localization and transport experiments, a synthetic and codon-optimized DNA fragment (GenScript, Leiden, Netherlands) referring to a hybrid *MtABCG40* sequence, consisting of 1044 bp of gDNA and 3741 bp of cDNA, was cloned into pMDC43 vector between the *Sgs*I (*Asc*I) and *Pac*I restriction sites (GenScript) (Curtis and Grossniklaus 2003). ATPase-deficient version of MtABCG40 (MtABCG40^-ATPase^ - E344Q and E1029Q) was generated by GenScript.

For RNAi silencing, a 188 bp cDNA fragment of *MtABCG40* 5’UTR or a 200 bp cDNA fragment of *MtLOG3* 3’UTR were amplified with KOD Hot Start DNA Polymerase (Novagen) and cloned into pDONR/Zeo vector (Thermo Fisher) using Gateway Recombination Cloning Technology. The obtained entry clone was subsequently recombined with pK7GWIWG2(II):DsRED binary vector (Limpens et al., 2005). A list of primers can be found in Table S1C.

Composite plants with transgenic roots were obtained through *Agrobacterium rhizogenes* Arqua1-mediated transformation, described by (Boisson-Dernier et al., 2001) with modifications. Shortly, the plants after transformation were kept at 20°C on full-strength Fähraeus agar plates containing 1 mM aminoisobutyric acid. After a week, emerged and potentially non-transgenic hairy roots were removed and the plants were transferred onto fresh Fähraeus, full-strength or 0 mM NH_4_NO_3_, at 23°C for 2-3 weeks to obtain transgenic roots. The identification of transgenic roots was possible due to an antibiotic or DsRed selection, depending on the vector used.

### MtABCG40 transport assays

For protoplast transport assays, protoplasts were prepared from Agrobacterium- transfected *N. benthamiana* leaves and [^14^C]-*t*Z and [^3^H]-IAA export was quantified as described previously (Henrichs et al., 2012). In short, tobacco mesophyll protoplasts were prepared 4 days after Agrobacterium-mediated transfection with *35S:GFP-MtABCG40* or *35S:GFP-MtABCG40^-ATPase^* or the empty vector control. Equal protoplast loading was achieved by diffusion on ice and export was determined by separating protoplasts and supernatants by silicon oil centrifugation. Relative export from protoplasts was calculated from exported radioactivity into the supernatant as follows: (radioactivity in the supernatant at time t = x min) - (radioactivity in the supernatant at time t = 0)) * (100%)/ (radioactivity in the supernatant at t = 0 min). In some cases, relative import into tobacco protoplasts was determined by separating protoplasts and supernatants by silicon oil centrifugation. [^14^C]-*t*Z loading was calculated from imported radioactivity into protoplasts as follows: (radioactivity in the protoplasts at time t = x min) - (radioactivity in the protoplasts at time t = 0)) * (100%)/ (radioactivity in the protoplasts at t = 0 min); presented are mean values from > 4 independent transfections and protoplasts preparations.

### Microscopic observations and staining

For subcellular localization experiments, *35S:GFP-MtABCG40*, *35S:GFP-MtABCG40^-ATPase^, 35S:ABCB1-RFP* (Henrichs et al., 2012) or *35S:HDEL-mCherry* (Liang et al., 2015) were expressed in *Nicotiana benthamiana* leaf tissues by *Agrobacterium tumefaciens*-mediated leaf infiltration as described previously (Henrichs et al., 2012). For confocal laser scanning microscopy, a Leica SP8 confocal laser microscope was used and confocal settings were set to record the emission of GFP (excitation 488 nm, emission 500–550 nm) and RFP/mCherry (excitation 543 nm, emission 580-640 nm).

Transgenic roots carrying a GUS (*β-glucuronidase*) reporter gene were stained using 5- bromo-4-chloro-3-indolyl-β-D-glucuronide (Gallagher, 1992) with an addition of 20% methanol (Kosugi et al., 1990). *M. truncatula* roots inoculated with *S. meliloti* 1021 (pXLGD4) strain were stained using 5-bromo-6-chloro-3-indolyl-β-d-galactopyranoside (Magenta-Gal, Sigma-Aldrich), according to the protocol described previously by Jarzyniak et al. (2021).

For measurement of RAM size, roots of 10-day-old plants grown on 0 or 1 mM NH_4_NO_3_ Fähraeus agar plates were collected. The length of RAMs was measured as a distance between a stem cell niche and the elongation/differentiation zone (EDZ), indicated by an onset of elongation of the second layer of cortical cells (Shen et al., 2019). The cells were visualized under Leica TCS SP5 laser scanning confocal microscope using Nomarski optics. The material was analyzed in two independent experiments, with more than 14 RAMs collected per one repetition.

For analysis of the pace of cell divisions within 12-hour lateral root and nodule primordia, bent root fragments or spot-inoculated root fragments, respectively were stained with EdU (5-ethynyl-2′-deoxyuridine; Thermo Fisher) and modified pseudo-Schiff propidium iodide (PI) according to the protocol described by (Schiessl et al., 2019). The samples were subsequently cleared by incubation in chloral hydrate (Sigma-Aldrich) solution and mounted in Hoyer’s medium according to the protocol of (Truernit et al., 2008). Microscopic observations were carried out using Leica TCS SP5 laser scanning confocal microscope with an excitation filter set at 488 nm and emission filters set to 572–625 nm and 505–600 nm for propidium iodide and EdU, respectively. Finally, the number of EdU-stained nuclei in each of the analyzed root fragments per *mtabcg40-1* and respective WT plants from two independent biological repeats was counted.

### Gene expression analyses

The isolation of total RNA from the collected samples was performed with the use of the RNeasy Mini Kit (QIAGEN, Hilden, Germany) according to the manufacturer’s instructions. Removal of the DNA from the samples was carried out through a DNase I treatment (QIAGEN). The cDNA was then synthesized with an Omniscript Reverse Transcription (RT) Kit (QIAGEN). Quantitative PCR reactions were carried out in a CFX Connect Real-time PCR detection system (BioRad, Hercules, California, USA) using iTaq Universal SYBR Green Supermix (BioRad) with at least three biological replicates each with three technical repeats. The relative mRNA expression levels were normalized to *Mtactin* and calculated using the ΔΔCt method. A list of primers can be found in Table S1D.

The heat map data were generated by dividing a relative (normalized to *Mtactin*) expression level of a gene in a specific NH_4_NO_3_ concentration by its expression level in 0.5 mM NH_4_NO_3_, resulting in a fold change (FC) value. Obtained FC value was then expressed logarithmically, resulting in a logFC value. The intensity of red color filling every cell on the graph reflects a specific logFC value, starting with the lowest (-0.18) and ending up with the highest one (0.70).

### Phytohormone quantification

For CK and auxin quantification, seeds of mutant (*mtabcg40-1, mtabcg40-2*) and corresponding WT plants after germination were transferred onto 0 mM NH_4_NO_3_ Fähraeus agar plates. 50 RAMs of 10-day-old seedlings per sample were collected into sterile high-purity water and lyophilized. Three or two biological replicates per mutant/WT were prepared. Endogenous levels of cytokinin metabolites and IAA were determined by LC-MS/MS methods (Svacinova et al., 2012; Pencik et al., 2018). 50 RAMs were homogenized and extracted in 0.5 ml of modified Bieleski buffer (60% MeOH, 10% HCOOH and 30% H_2_O) with added stable isotope-labeled internal standards (0.25 pmol of CK bases, ribosides, N-glucosides, 0.5 pmol of CK O-glucosides and nucleotides, and 5 pmol of [^13^C_6_] IAA per sample). Phytohormones were determined using an ultra-high performance liquid chromatography (Aquity UPLC® I-class system; Waters, Milford, MA, USA) with electrospray interface tandem mass spectrometry (Xevo TQ-S, Waters, Manchester, UK) using stable isotope-labelled internal standards as a reference.

### Statistical analyses

Statistical analyses were performed using the GraphPad Prism software (v.9.4.1). The normality assumption was verified on residuals by the usage of the Shapiro–Wilk and Kolmogorov-Smirnov normality test. The normal distribution of residuals from each group was analyzed separately. If the normality assumptions were met, parametric tests (i.e., two-tailed Student’s t-test, two-tailed Student’s t-test with Welch correction, one-way ANOVA with a post hoc Tukey’s multiple comparison test) were applied. If the normality assumptions were not met, non-parametric tests (i.e., two-tailed Mann–Whitney test, Kruskal-Wallis test with a post hoc Dunn’s multiple comparison test) were applied. Details regarding statistical analyses can be found in Excel File S1.

## Results

### Lateral root density is negatively regulated by MtABCG40 in response to environmental and internal cues

*MtABCG40* (*Medtr7g098300*) encodes a full-size ABC transporter that belongs to a legume-specific clade of the G subfamily (Banasiak and Jasiński, 2014; Jarzyniak et al., 2021). High expression of *MtABCG40* in the root prompted us to determine a possible role of the encoded transporter in this organ (Fig. 1A). Due to a reported multifaceted influence of environmental stimuli, such as nitrogen status, on root morphology, we investigated *MtABCG40* expression at different concentrations of ammonium nitrate (NH_4_NO_3_). Compared to plants grown on media with added NH_4_NO_3_, plants grown on solid media not supplemented with nitrogen (here referred to as 0 mM NH_4_NO_3_) exhibited the highest *MtABCG40* expression and the lowest lateral root density (Fig. 1B). The decline in the lateral root density, resulting from an enlargement in the spacing between adjacent lateral roots, was caused by an acceleration of primary root elongation and, to a lesser extent, a decrease in the lateral root number (Fig. S1A,B). We also investigated a possible relationship between *MtABCG40* expression and LR density upon treatment with hormones known to trigger changes in root architecture, auxin and CKs. *MtABCG40* mRNA abundance decreased after indole-3-acetic acid (IAA) treatment, which was associated with increased lateral root density; in contrast, application of 6-BAP, a synthetic CK, resulted in an induction of *MtABCG40* expression accompanied by a decline in lateral root density (Fig. 1C,D). For both phytohormone treatments, the changes in lateral root density resulted from the alteration in primary root length and lateral root number (Fig. S1C- F).

**Fig. 1.**
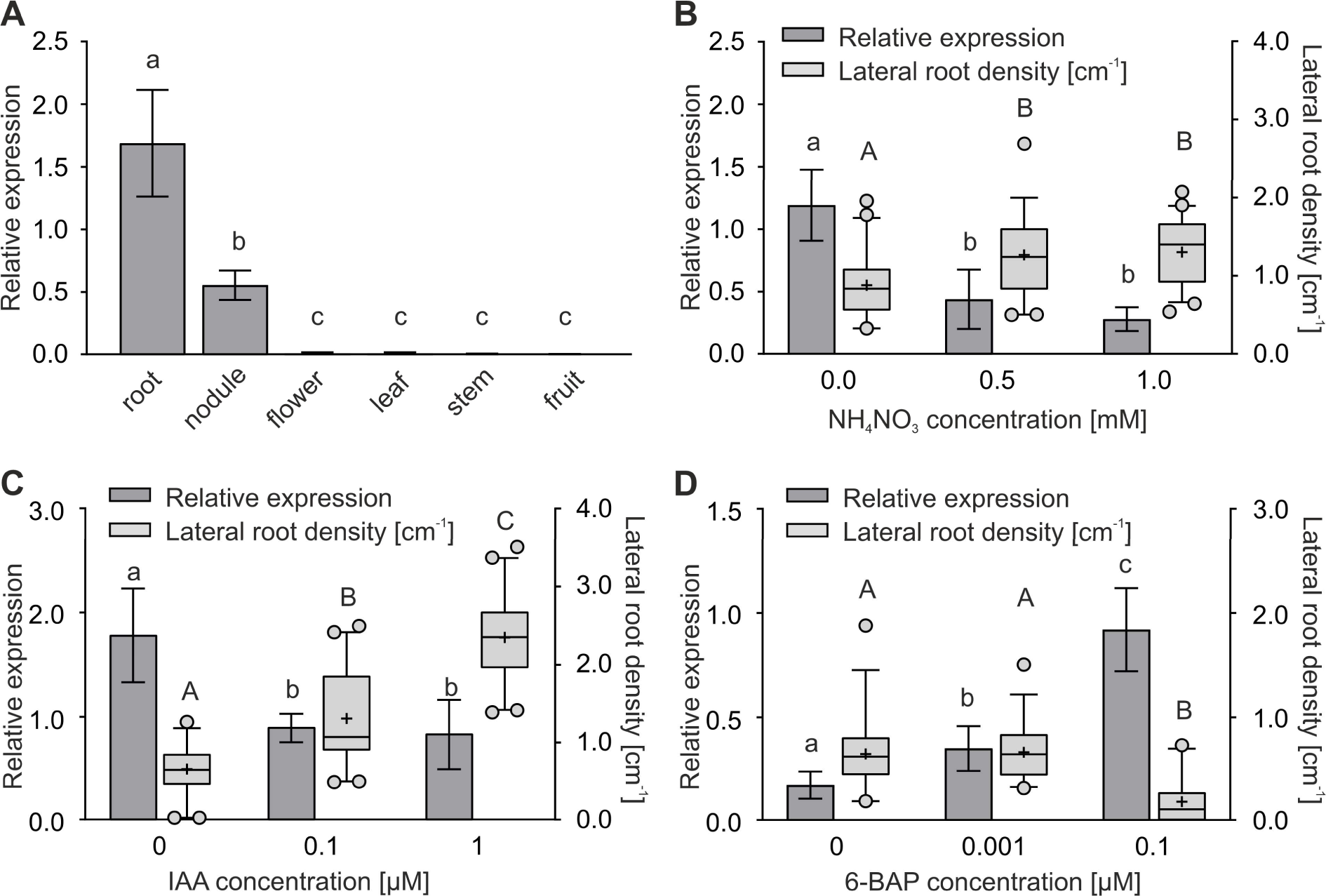
A negative relationship was found between *MtABCG40* expression and LR density. A, *MtABCG40* transcripts were detected only in roots and nodules. B-D, Expression of *MtABCG40* and lateral root density at different concentrations of ammonium nitrate (NH_4_NO_3_) (B), indole-3-acetic acid (IAA) (C), and 6-benzylaminopurine (6-BAP) (D). Transcript levels were measured by quantitative real-time PCR and normalized to *Mtactin*. Expression data represent the mean ± SD of three independent biological experiments and two or three technical repeats. Identical or different lowercase letters indicate no or significant differences in expression, respectively; *P* < 0.05 (A-D). Significant differences were determined by the one-way ANOVA with a post hoc Tukey’s multiple comparison test (A-D). The box plots present the lateral root density for 31-45 roots per condition obtained from three independent biological experiments. For each box-and-whiskers plot, the central black line represents the median; ‘+’ represents the mean; the box extends from the 25th to 75th percentiles; and the whiskers are drawn down to the 5th percentile and up to the 95th. Points below and above the whiskers are drawn as individual dots. Identical or different uppercase letters indicate no or significant differences in the lateral root density, respectively; *P* < 0.05 (B-D). Significant differences were determined by one-way ANOVA with a post hoc Tukey’s multiple comparison test (B) or Kruskal–Wallis test with a post hoc Dunn’s multiple comparison test (C and D).

Two *M. truncatula* lines with tobacco retrotransposon (*Tnt1*) insertions in the 22^nd^ exon (NF21323, *mtabcg40-1*) and 1^st^ intron (NF17891, *mtabcg40-2*) of *MtABCG40* were identified (Fig. S2). When grown on 0 mM NH_4_NO_3_ media, the mutants exhibited a significant increase in lateral root density in comparison to WT plants, which could indicate that *MtABCG40* negatively influences this trait (Fig. 2A). The phenotype resulted from a reduction in the primary root length (Fig. 2B) and an increase in the lateral root number (Fig. 2C). Notably, the observed differences were less pronounced when nitrogen was present (1 mM NH_4_NO_3_) (Fig. S3). Furthermore, silencing of *MtABCG40* expression also caused an increase in the lateral root density of transgenic hairy roots grown on media not supplemented with nitrogen (Fig. 2D-F).

**Fig. 2.**
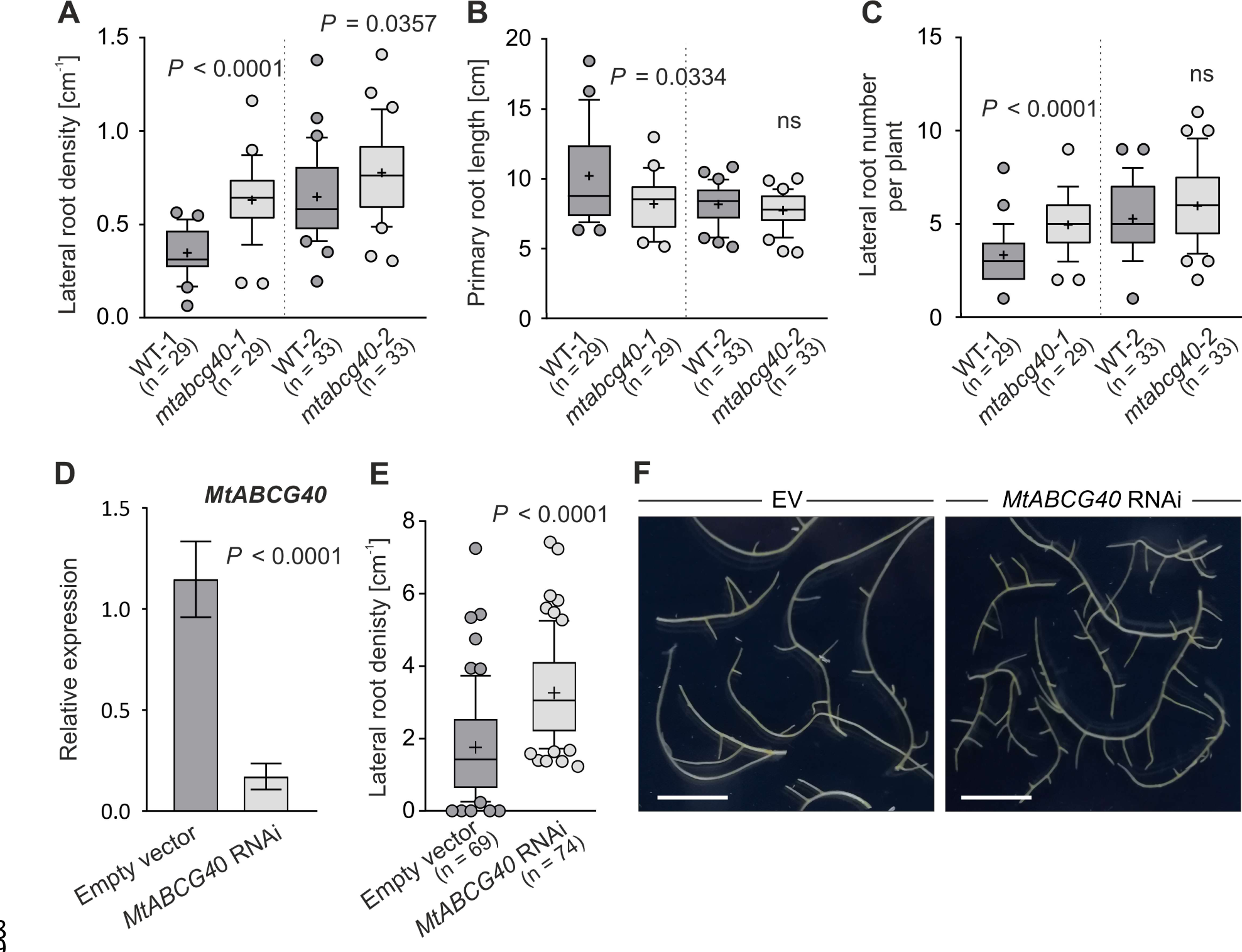
Effect of *MtABCG40* mutation or *MtABCG40* RNAi silencing on root morphology under nitrogen starvation. Lateral root density (A), primary root length (B), and lateral root number (C) in control (WT) and mutant (*mtabcg40*) plants. D, *MtABCG40* transcript accumulation in *MtABCG40* RNAi-silenced hairy roots in comparison to the empty vector (EV)-transformed control. Transcript levels were measured by quantitative real-time PCR and normalized to *Mtactin*. Expression data represent the mean ± SD of two independent biological experiments and three technical repeats. E, Lateral root density of *MtABCG40* RNAi-silenced hairy roots in comparison to the EV-transformed control. F, Images of *MtABCG40* RNAi-silenced hairy roots in comparison to the EV-transformed control. Scale bars: 1 cm. For each box-and-whiskers plot, the central black line represents the median; ‘+’ represents the mean; the box extends from the 25th to 75th percentiles; and the whiskers are drawn down to the 10th percentile and up to the 90th. Points below and above the whiskers are drawn as individual dots; n represents the number of individual plants obtained from three independent biological experiments (A-C and E). Significant differences from the control plants were determined by two-tailed Student’s t test with Welch correction (A and D), two-tailed Mann–Whitney test (B, C and E); ns, not significant.

### MtABCG40 is a *bona fide* plasma membrane cytokinin importer

Since 6-BAP treatment induced *MtABCG40* expression (Fig. 1D) and given MtABCG40’s close phylogenetic relationship to the previously described CK transporter MtABCG56 (Fig. S4) (Jarzyniak et al., 2021), we decided to study the possible role of MtABCG40 in CK translocation. Initially, in addition to 6-BAP, we tested the effect of other CKs on *MtABCG40* expression and observed that the biologically active CKs, iP and *t*Z, induced *MtABCG40* mRNA accumulation in roots (Fig. 3A,B).

**Fig. 3.**
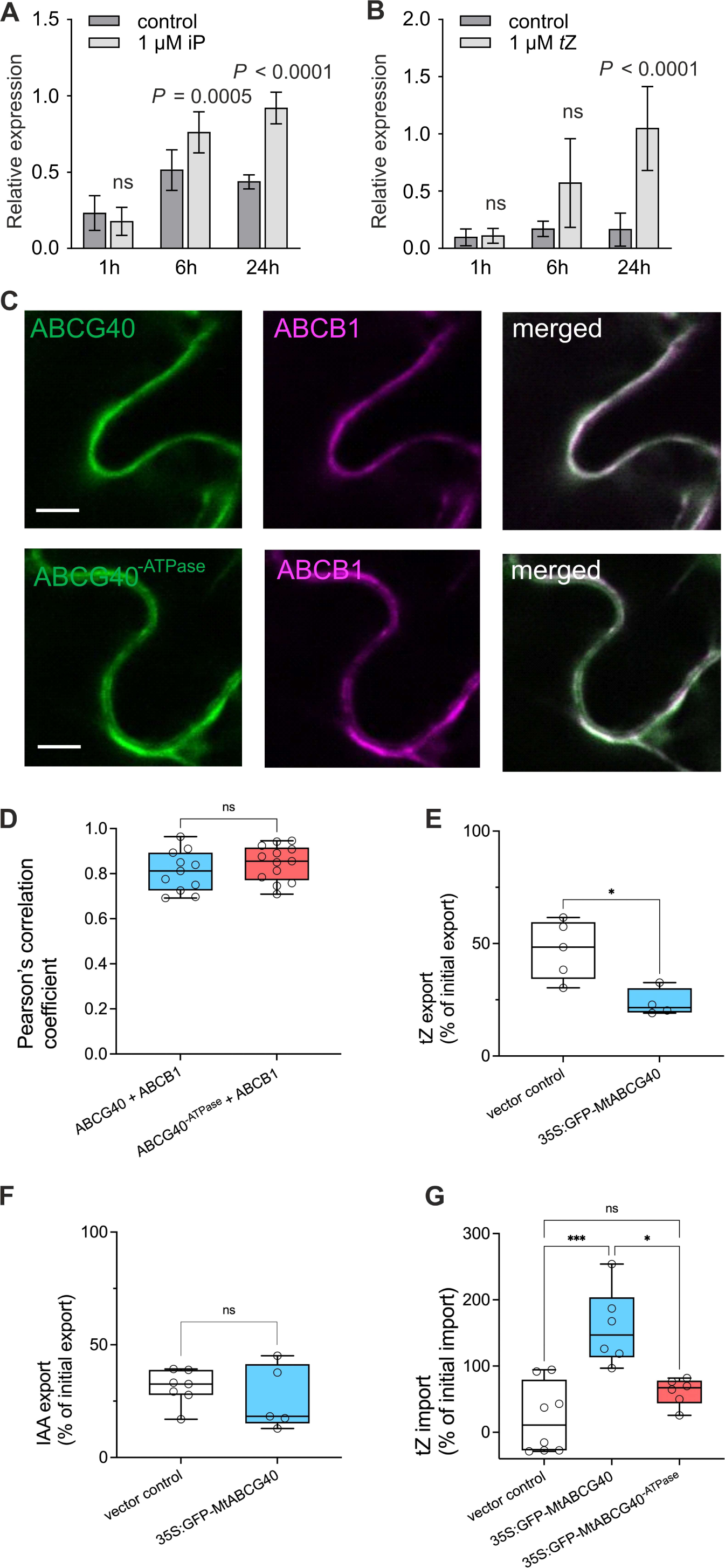
MtABCG40 is a *bona fide* plasma membrane (PM) cytokinin (CK) importer. A and B, *MtABCG40* expression profile upon CK treatment. Transcript levels of roots mock-treated or treated with 1 μM isopentenyladenine (iP) (A) and 1 μM *trans-*zeatin (*t*Z) (B), measured by quantitative real-time PCR and normalized to *Mtactin* from *Medicago truncatula*. Data represent the mean ± SD of three independent biological experiments and three technical repeats. Significant differences from the mock- treated plants determined by two-tailed Mann–Whitney test (A and B). C and D, PM localization of the GFP-MtABCG40 and GFP-MtABCG40^-ATPase^ fusion proteins in Agrobacterium-infiltrated *Nicotiana benthamiana* leaf epidermal cells with the PM marker, ABCB1 (C). Pearson’s correlation coefficient between GFP-MtABCG40 and ABCB1-RFP and between GFP-MtABCG40^-ATPase^ and ABCB1-RFP (D). GFP-MtABCG40, GFP-MtABCG40^-ATPase^ and ABCB1-RFP images pseudo-coloured in green and magenta, respectively. Scale bars, 50 μm. E-G, Transport experiments in tobacco protoplasts expressing *GFP-MtABCG40* and *GFP*-*MtABCG40^-ATPase^*. Relative [^14^C]-*t*Z (E) and [^3^H]-indole-3-acetic acid (IAA) (F) export from tobacco protoplasts, as well as [^14^C]-*t*Z import experiments into tobacco protoplasts (G). Data represent the result from a minimum of 4 independent experiments (transfections and protoplast preparations). Significant differences from the vector control were determined by two-tailed Student’s t test (E-F) and one-way ANOVA with a post hoc Tukey’s multiple comparison test (G) are indicated as follows: *P < 0.05, **P < 0.01, ****P < 0.0001; ns, not significant. Time kinetics data can be found in Fig. S5.

To demonstrate that MtABCG40 is directly involved in the translocation of CKs, transport experiments with radiolabelled *t*Z were carried out. First, we expressed *GFP- MtABCG40* under the control of the *CaM35S* promoter in tobacco using Agrobacterium-mediated leaf infiltration. Colocalization (Pearson’s correlation coefficient (PCC) = 0.810 +/- 0.027) with the well-established PM marker AtABCB1 (Geisler et al., 2005, Henrichs et al., 2012) indicated a PM localization of MtABCG40 (Fig. 3C,D). However, we also found partial colocalization with the ER marker HDEL-mCherry (Liang et al., 2015) (PCC = 0.680 +/- 0.026; Fig. S5A,B). A higher correlation coefficient for PM than ER locations indicates that the main portion of MtABCG40 resides on the PM, while a smaller portion resides on the ER.

Subsequently, protoplasts isolated from the transformed leaves were loaded with [^14^C]- *t*Z, and *t*Z export was determined by separating protoplasts from supernatants. Compared to the vector control, protoplasts expressing *GFP-MtABCG40* under the *CaM35S* promoter exported significantly less [^14^C]-*t*Z (Figs 3E, S5C). [^14^C]-*t*Z export catalysed by GFP-MtABCG40 was specific, as the diffusion control auxin (IAA), which is transported by some members of the ABCB family (Geisler et al., 2005), was not significantly affected by the expression of *GFP- MtABCG40* (Figs 3F, S5D). Reduced [^14^C]-*t*Z export together with a predominant PM localization suggests that GFP-MtABCG40 plays a role in the net import of CKs. To further foster this role in CK import, we conducted [^14^C]-*t*Z uptake experiments in protoplasts expressing *GFP-MtABCG40*. As expected, *GFP-MtABCG40* protoplasts imported significantly more [^14^C]-*t*Z than the vector control (Figs 3G, S5E).

To verify that *t*Z import by MtABCG40 was direct and ATP-dependent and that upregulation of tobacco endogenous uptake systems did not cause the increased uptake, we investigated the behaviour of an ATPase-deficient version of MtABCG40 (MtABCG40^-ATPase^) by mutating E344Q and E1029Q, conserved key residues in the Walker motifs. MtABCG40^-ATPase^ was likewise predominantly expressed on the PM (Fig. 3C,D; PCC = 0.846 +/- 0.022) and partially on the ER (Fig. S5A,B; PCC = 0.643 +/- 0.020). Importantly, *t*Z imported by MtABCG40^-ATPase^ was not significantly different from the vector control (Figs 3G, S5E).

### MtLOG3 activity affects Medicago root morphology upon nitrogen depletion

Active CKs are formed when sugar components are cleaved from their monophosphate riboside forms by LOG enzymes (Kurakawa et al., 2007). Therefore, we investigated a possible relationship between *MtABCG40*, *MtLOG* and *MtLOG-like* genes under various NH_4_NO_3_ concentrations. We found that five of these genes, namely, *MtLOG2* (*Medtr1g064260*), *MtLOG3* (*Medtr1g057020*), *MtLOG-like 1* (*Medtr1g015830*), *MtLOG-like 3* (*Medtr3g113710*) and *MtLOG-like 4* (*Medtr4g058740*) (Mortier et al., 2014; van Zeijl et al., 2015), showed increased expression under nitrogen depletion, which also positively influenced the level of *MtABCG40* mRNA (Fig. 4A). We then focused on the *MtLOG3* (van Zeijl et al., 2015), which is the closest homologue of Arabidopsis *LOG7* (*AT5G06300*), a nitrogen starvation-induced gene with expression in lateral root-forming pericycle cells (Fig. S6) (Bargmann et al., 2013; Walker et al., 2017). Silencing of *MtLOG3* expression caused an increase in lateral root density (Fig. 4B,C), similar to what was observed for *MtABCG40* RNAi-silenced roots (Fig. 2D-F) and *mtabcg40* mutants (Fig. 2A). This observation prompted us to investigate the *MtABCG40* and *MtLOG3* spatial expression patterns in roots. To do this, we generated *M. truncatula* composite plants that contained *MtABCG40* and *MtLOG3* promoters that drive the expression of either nucleus-localized versions of green fluorescent protein (*ProMtABCG40:NLS-GFP*) or tdTomato (*ProMtLOG3:NLS-tdTomato*) and *β-glucuronidase* (*ProMtABCG40:GUS* and *ProMtLOG3:GUS*) reporter genes. Transgenic roots subjected to nitrogen deficiency for three weeks revealed that *MtABCG40* is expressed mainly in the root stele, a site of radial, short-distance CK translocation (Ko et al., 2014; Aubry et al., 2019). A reporter signal was also observed in the pericycle and endodermis, layers known to be involved in initial cell divisions during lateral root primordium formation in *M. truncatula* (Herrbach et al., 2014). Additionally, *MtABCG40* expression was observed in the RAM, in which CKs influence meristem size and consequent organ elongation (Dello Ioio et al., 2007) (Fig. 4D). Under the same conditions, *MtLOG3* expression overlapped with that of *MtABCG40* in the root stele, but unlike *MtABCG40, MtLOG3* was expressed in the root cap, a site of CK biosynthesis (Aloni et al., 2005) (Fig. 4E).

**Fig. 4.**
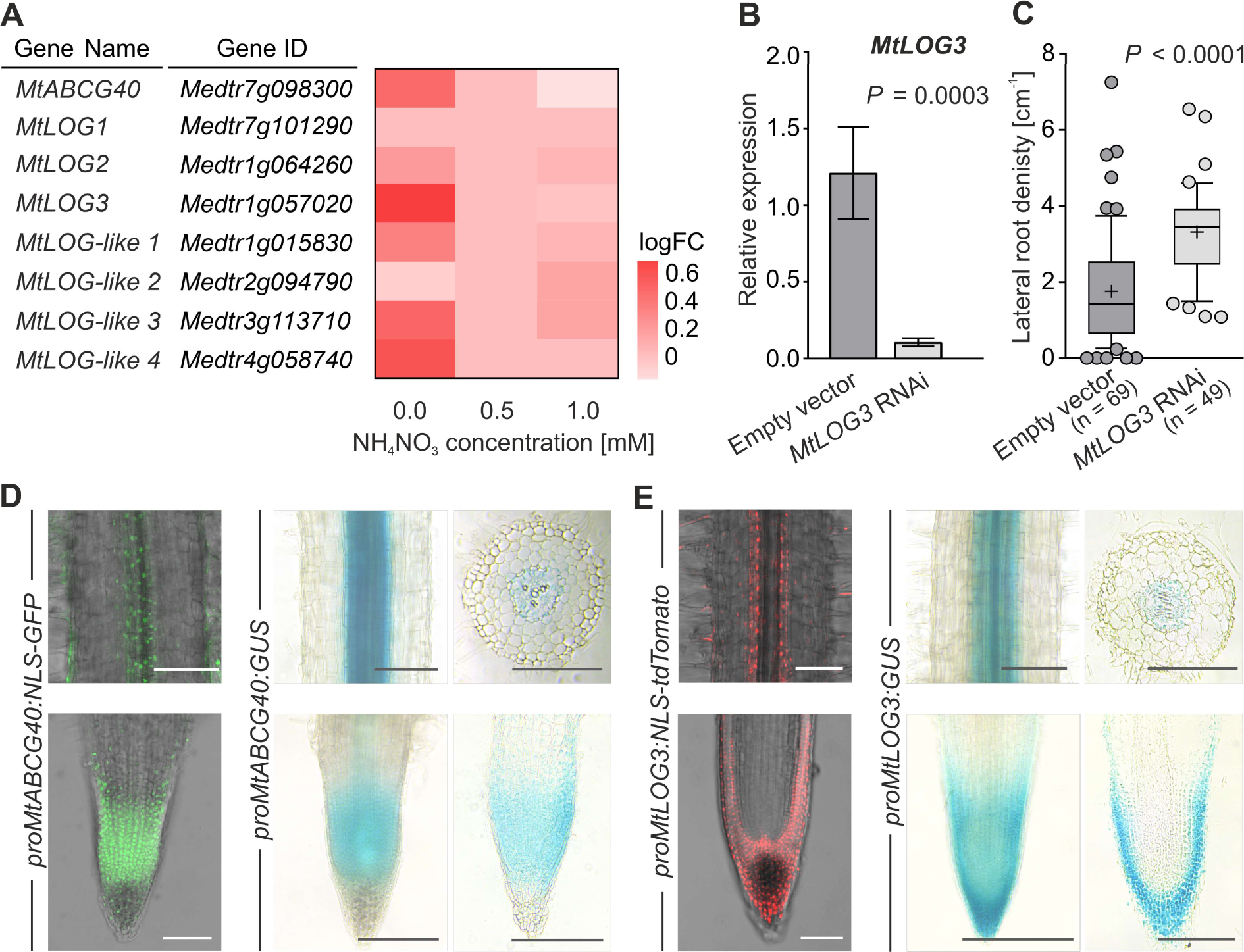
*MtLOG3* shares the expression profile in the root with *MtABCG40* and regulates root architecture upon nitrogen starvation. A, Heatmap showing the expression of *MtABCG40*, *MtLOG*, and *MtLOG-like* genes in root samples across NH_4_NO_3_ concentrations. Expression is shown as log2-fold changes. B, *MtLOG3* transcript accumulation in *MtLOG3* RNAi-silenced hairy roots in comparison to the empty vector (EV)-transformed control. Transcript levels were measured by quantitative real-time PCR and normalized to *Mtactin*. Expression data represent the mean ± SD of two independent biological experiments and three technical repeats. Significant differences from the control plants were determined by two-tailed Student’s t test with Welch correction. C, Lateral root density of *MtLOG3* RNAi-silenced hairy roots in comparison to the empty vector (EV)-transformed control. For each box-and-whiskers plot, the central black line represents the median; ‘+’ represents the mean; the box extends from the 25th to 75th percentiles; and the whiskers are drawn down to the 10th percentile and up to the 90th. Points below and above the whiskers are drawn as individual dots. Significant differences from the EV-transformed control were determined by a two-tailed Mann–Whitney test; n represents the number of individual roots obtained from three independent biological experiments. D-E, Tissue-specific expression of *MtABCG40* (D) and *MtLOG3* (E) in transgenic roots carrying the *proMtABCG40:NLS-GFP, proMtABCG40:GUS*, *proMtLOG3:NLS-tdTomato*, and *proMtLOG3:GUS* constructs, respectively. Each panel contains images from confocal microscopy (left side) and light microscopy with intact roots and cross-sections (right side), including the root apical meristem (RAM). The images are representative of n > 20 roots from three independent experiments (transformations). Scale bars, 100 μm.

### Cytokinin response in the root apical meristem is inhibited by MtABCG40

It is well documented that CKs decrease the size of RAM (Dello Ioio et al., 2007; Dello Ioio et al., 2008). To explain the changes in root length between WT and *mtabcg40* plants (Fig. 2B), we analysed the sizes of their RAMs as the distance between the quiescent centre (QC) and the elongation/differentiation zone (EDZ). Compared to WT, the *mtabcg40* mutants exhibited shorter RAMs under nitrogen deficiency (Fig. 5A,B). This phenotype of *mtabcg40* was indistinguishable from that of WT when plants grew on medium supplemented with 1 mM NH_4_NO_3_ (Supporting Fig. S7). Additionally, the negative role of MtABCG40 in CK responses upon nitrogen starvation was demonstrated by the higher expression of *MtRR4*, a type-A CK response regulator (Gonzalez-Rizzo et al., 2006), in the root tip of *mtabcg40-1* compared to the WT (Fig. 5C,D). Notably, no difference was observed in the overall active CK content within those RAMs (Fig. 5E), suggesting that the distribution of active CKs rather than their total concentration is affected by loss of *MtABCG40*. As CKs often antagonize auxin outcomes in meristematic regions, we measured the amount of auxin (IAA) in the RAMs of *mtabcg40* lines and observed an increased IAA level compared to the WT (Fig. 5F). Moreover, we observed that the *DR5* auxin-response reporter in RAMs of *MtABCG40* RNAi-silenced roots was upregulated (Fig. 5G).

**Fig. 5.**
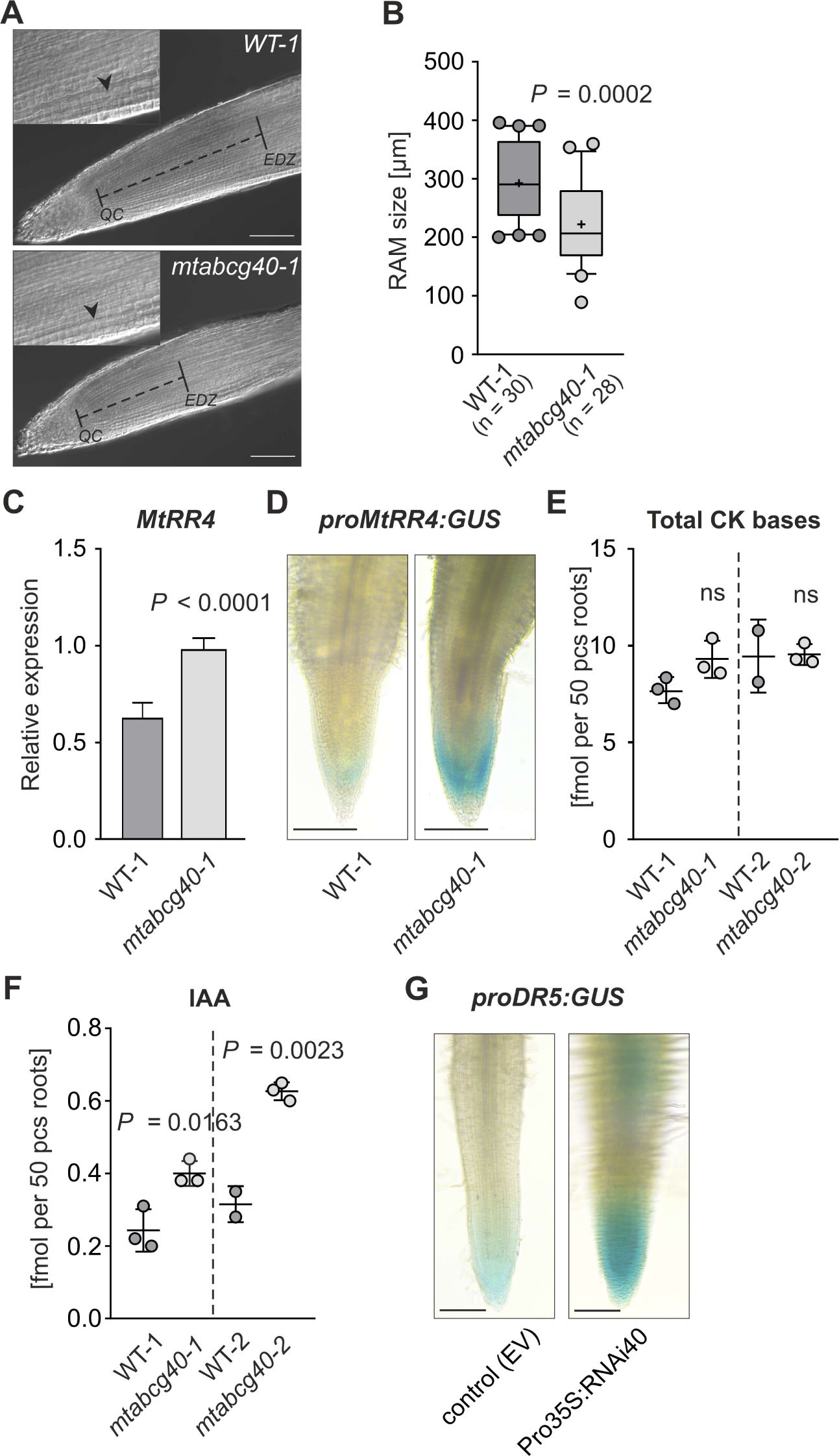
Mutation of *MtABCG40* reduces the size of the *Medicago truncatula* root apical meristem (RAM) upon nitrogen deficiency (0 mM NH_4_NO_3_). A and B, Comparison of the size of RAM between WT-1 and *mtabcg40-1*. Nomarski image of the RAM and apical region in the elongation/differentiation zone (EDZ) of a primary root 10 days after germination. The size of the RAM (black dotted line) was determined as the distance from the quiescent centre (QC) to the EDZ. The apical border of the EDZ was defined by the first elongated cortical cell of the second cortical layer (arrowhead) (A). Graph showing the RAM size of WT-1 and the *mtabcg40-1* mutant. For each box-and-whiskers plot, the central black line represents the median; ‘+’ represents the mean; the box extends from the 25th to 75th percentiles; and the whiskers are drawn down to the 10th percentile and up to the 90th. Points below and above the whiskers are drawn as individual dots; n represents the number of analysed RAMs (B). C-E, Assessment of the level of CK response and cytokinin (CK) bases (*trans-*zeatin, *cis-*zeatin, isopentenyladenine, dihydrozeatin) in RAMs of WT-1 and *mtabcg40-1* mutant. Expression analysis of *MtRR4*, a type-A response regulator, in WT-1 and *mtabcg40-1*. Transcript levels were measured by quantitative real-time PCR and normalized to *Mtactin* (C). *ProMtRR4:GUS* activity in corresponding WT and *mtabcg40-1* roots. Images are representative of n > 10 roots. Scale bars, 100 μm (D). Total CK bases concentration in WT-1 and *mtabcg40-1* (E). F and G, Assessment of auxin (indole-3-acetic acid, IAA) levels and responses in RAMs of WT-1 and the *mtabcg40-1* mutant. IAA concentration in WT-1 and *mtabcg40-1* (F). *DR5:GUS* reporter expression in *MtABCG40* RNAi-silenced and control (EV) roots; representative images from 10 roots of each construct. Scale bars, 100 μm (G). Data represent the mean ± SD of two independent biological experiments (B), three independent biological experiments and three technical repeats (C), and two or three biological replicates (E and F). Significant differences from the control plants determined by two-tailed Mann–Whitney test (B), two-tailed Student’s t test (C, E and F); ns, not significant.

### Lateral root initiation is enhanced in *mtabcg40* plants

To further explore the increased lateral root number in *mtabcg40* plants (Fig. 2C), we used gravitropic stimulation of lateral root initiation on 0.5 mM NH_4_NO_3_ (Fig. S8) combined with EdU (5-ethynyl-2′-deoxyuridine)-staining of DNA in dividing cells (Schiessl et al., 2019). In lateral root primordia that formed on 12-hour gravistimulated roots, a higher number of cells were observed in *mtabcg40-1* plants than WT-1 plants (Fig. 6A,B). Additionally, stronger expression of *MtABCG40* was observed in some particular cells on the margins of the root stele, most likely in the pericycle (Fig. 6C). Since CKs antagonize auxin-dependent initiation of lateral root formation (Laplaze et al., 2007), we assessed a possible effect of *MtABCG40* on the auxin response. By using once again gravitropic stimulation of lateral root initiation, we assessed the transcript level of the auxin-inducible gene *LATERAL ORGAN BOUNDARIES DOMAIN16* (*MtLBD16*), which marks the onset of lateral root organogenesis (Schiessl et al., 2019). We found that *MtLBD16* mRNA accumulation and the number of EdU-stained nuclei were enhanced in *mtabcg40* roots compared to WT roots in 0.5 mM but not 1 mM NH_4_NO_3_ (Figs 6, S9).

**Fig. 6.**
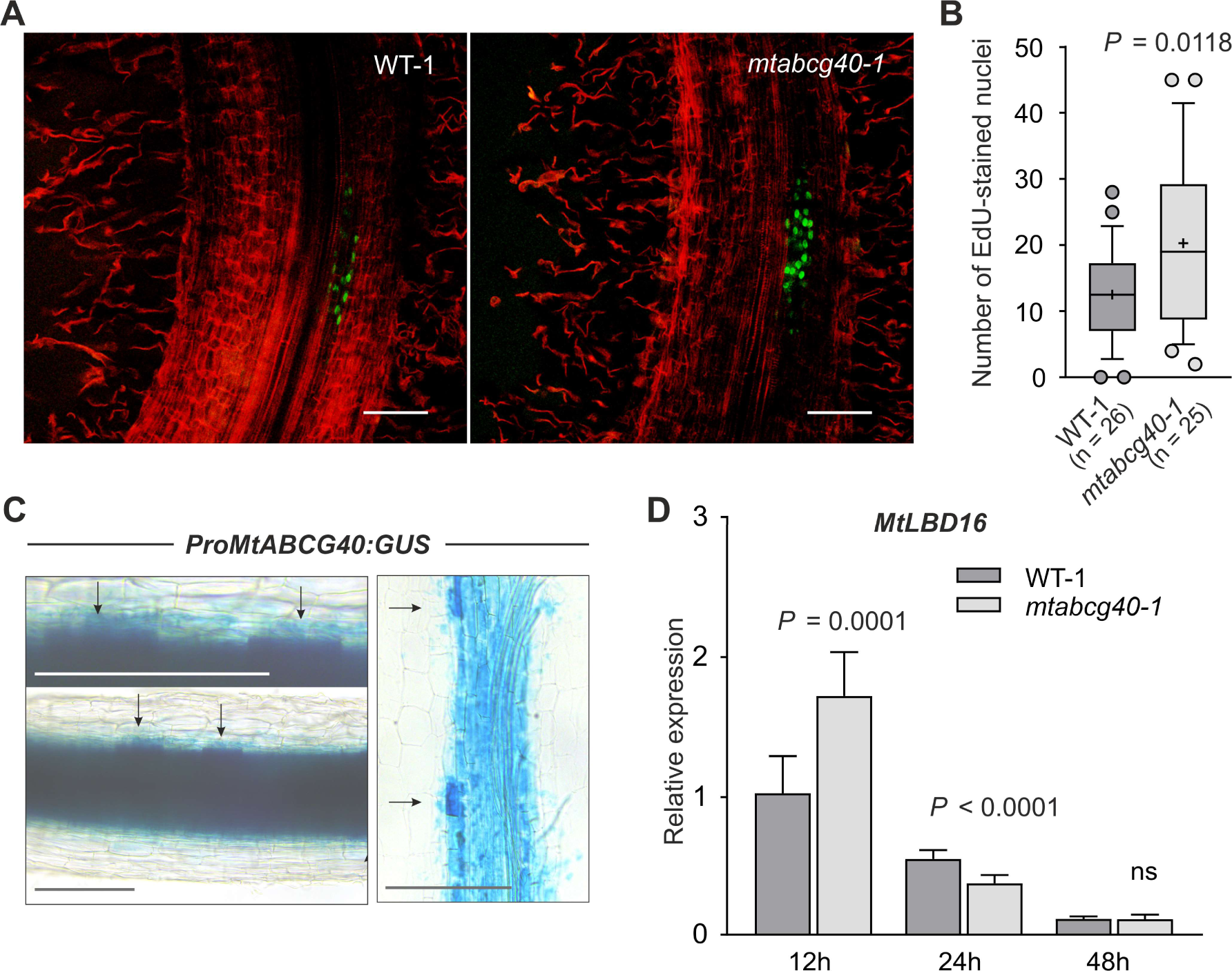
Lack of *MtABCG40* accelerates lateral root initiation in 0.5 mM NH_4_NO_3_. A and B, Comparison of the cell division rate within the lateral root primordium 12 hours postinduction (hpi) by gravitropic stimulation between WT-1 and *mtabcg40-1* mutant plants. Optical sections of representative lateral root primordia. Red propidium iodide demarks cell walls, and green EdU-labelled nuclei indicate DNA replication. Scale bars, 100 μm (A). Comparison of the number of EdU-stained nuclei within the lateral root primordium between WT-1 and *mtabcg40-1*; n represents the number of individual plants obtained from two independent experiments; for each box-and-whiskers plot, the central black line represents the median; ‘+’ represents the mean; the box extends from the 25th to 75th percentiles; and the whiskers are drawn down to the 10th percentile and up to the 90th. Points below and above the whiskers are drawn as individual dots (B). C, *ProMtABCG40:GUS* activity in root stele and the particular cells on the margins of the root stele (marked by arrows). Images are representative of n > 10 roots obtained from two independent transformations. Scale bars, 100 μm. D, Expression analysis of *MtLBD16* in *mtabcg40- 1* and WT-1 within the gravitropically stimulated bent fragments (2-3 mm) at the indicated time points, 12, 24 and 48 hpi. Transcript levels were measured by quantitative real-time PCR and normalized to *Mtactin*; data represent the mean ± SD of three independent biological experiments and three technical repeats. Significant differences from the WT plants determined by two-tailed Student’s t test with Welch correction (B), two-tailed Student’s t test (D); ns, not significant.

### Nodule initiation is enhanced in *mtabcg40* plants

Since lateral roots and nodules share overlapping developmental programs (Schiessl et al., 2019), we also examined the influence of MtABCG40 on nodulation. We initially assessed the response of *MtABCG40* to Nod factors (NFs). The latter are produced by the bacterial microsymbiont as a part of a chemical dialogue with plant roots in nitrogen-deprived conditions, which initiates nodule formation (Zuanazzi et al., 1998). We inoculated the roots of *M. truncatula* with suspensions of the following strains of symbiotic bacteria: *S. meliloti* E65, which constitutively overproduces NFs, and *S. meliloti* SL44, which cannot produce NFs, as a control. We revealed that *MtABCG40* was induced 24 h after inoculation with *S. meliloti* E65 (Fig. 7A). Interestingly, bacterial inoculation did not induce *MtABCG40* systemically, as it was observed for *TOO MUCH LOVE* (*TML*) gene, encoding a Kelch Repeat-Containing F-box Protein that is an inhibitor of nodule organogenesis in response to shoot-derived signals (i.a. cytokinins) during autoregulation of nodulation (AON) (Fig. S10) (Takahara et al., 2013; Gautrat et al., 2019). To quantify the effect of MtABCG40 on nodulation, we counted the nodule number of *mtabcg40* plants. Both mutant lines (*mtabcg40-1* and *mtabcg40-2*) produced more nodules than their respective wild types, implying that MtABCG40 plays a negative role in nodule formation (Fig. 7B). To determine whether the transporter affects the initiation of nodule primordia, we performed bacterial spot inoculation, in which a droplet of *S. meliloti* suspension was applied onto the susceptible zone (Fig. 7C). We detected that expression of *MtLBD16* was enhanced in *mtabcg40-1* compared to WT-1 plants within the spot-inoculated region (Fig. 7D). This finding was associated with a higher number of cells labelled with EdU in the *mtabcg40-1* nodule primordia in comparison to WT-1 (Fig. 7E,F). These results imply that MtABCG40 suppresses initial cell divisions in nodule formation.

**Fig. 7.**
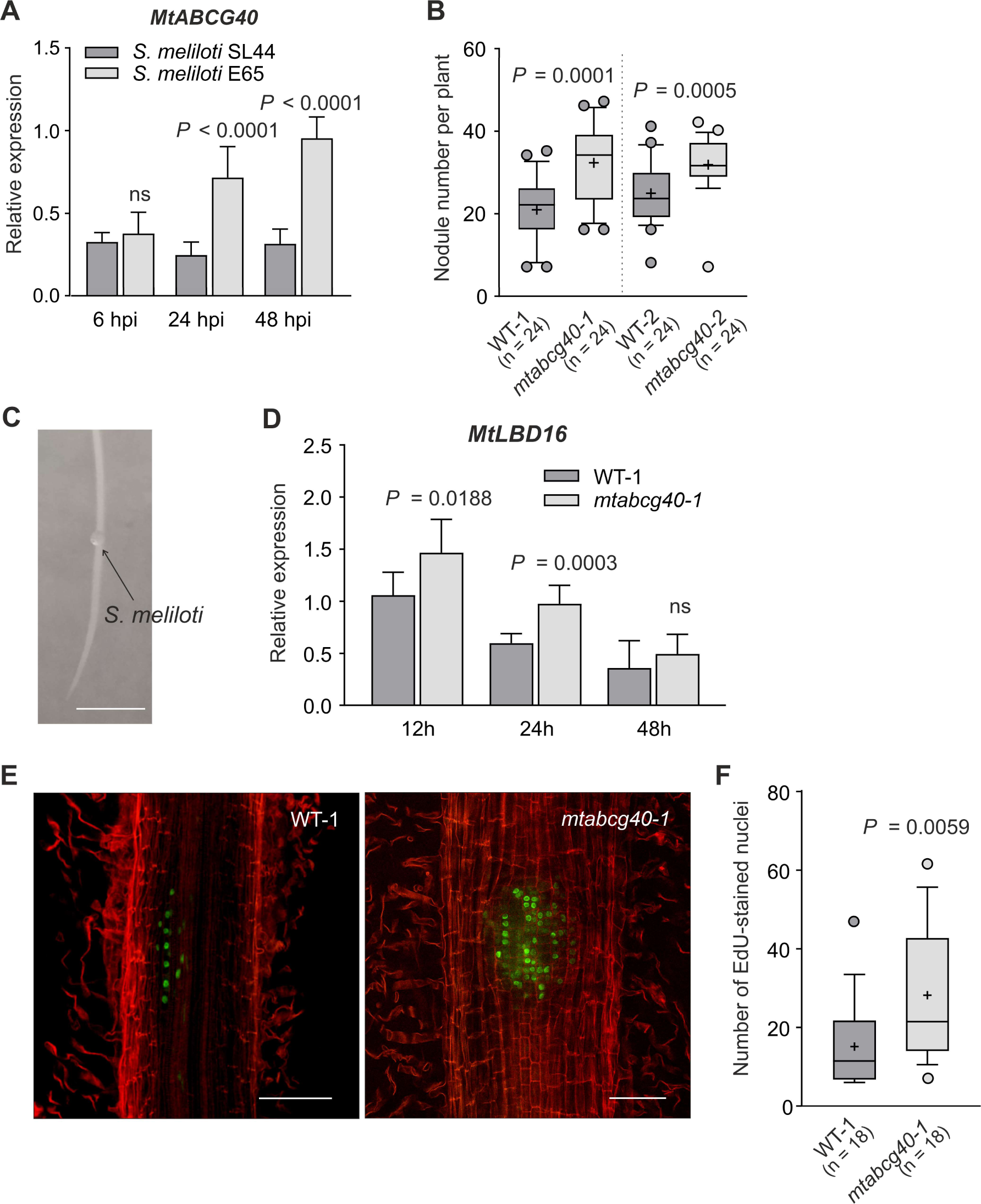
MtABCG40 negatively influences nodule number. A, Comparison of *MtABCG40* expression in roots flood-inoculated with suspensions of *Sinorhizobium meliloti* E65 (constitutively overproducing Nod factors) and SL44 (which cannot produce NFs). B, Nodule number formed on control (WT) and mutant (*mtabcg40*) plants 21 days after *S. meliloti* 1021 wild-type strain inoculation. C-F, Analysis of the auxin response and cell division rate in spot-inoculated fragments during the early stages of nodulation. A photo of root spot-inoculated with a drop of *S. meliloti* suspension. Scale bar: 0.5 cm (C). Expression analysis of *MtLBD16* in *mtabcg40* and WT-1 roots (D). E and F, Comparison of the cell division rate within nodule primordia between *mtabcg40-1* and WT-1. Optical sections of nodule primordia 12 hours post inoculation (hpi). Red propidium iodide demarks cell walls, and green EdU-labelled nuclei indicate DNA replication (E). Comparison of the number of EdU-stained nuclei within the nodule primordium between WT-1 and the *mtabcg40-1* mutant (F). Transcript levels were measured by quantitative real-time PCR and normalized to *Mtactin*; data represent the mean ± SD of three independent biological experiments and three technical repeats (A and D). n represents the number of individual plants obtained from three (B) or two (F) independent experiments; for each box-and-whiskers plot, the central black line represents the median; ‘+’ represents the mean; the box extends from the 25th to 75th percentiles; and the whiskers are drawn down to the 10th percentile and up to the 90th. Points below and above the whiskers are drawn as individual dots (B and F). Significant differences from the control plants determined by two-tailed Mann–Whitney test (A, B, D and F).

## Discussion

Root development is a postembryonic process that is highly flexible in response to fluctuations of nutrients in the environment. The change in root morphology is tightly linked to the activity of RAM and the formation of lateral roots, which are controlled by hormonal crosstalk (Jia and von Wiren, 2020). Our studies suggest that under nitrogen shortage, MtABCG40 is involved in the negative control of lateral root density (Figs 1B, 2A). The latter results from an enhanced elongation of the primary root and a decrease in lateral root number (Fig. 2B,C).

Earlier studies suggested that CK metabolism and translocation are modulated by nitrogen status (Takei et al., 2002). Notably, root cap-derived CKs inhibit RAM activity in N-sufficient conditions (Tsugeki and Fedoroff, 1999; Aloni et al., 2005). We found that nitrogen shortage leads to locally increased expression of *LOG* genes in the root (Fig. 4A), which is expected to promote the conversion of inactive CKs in the root cap into their free, biologically active forms (Fig. 4E) (Kurakawa et al., 2007). The lack of an inhibitory effect of the root cap-derived CKs on RAM activity and overall root elongation under N deficiency suggests that a mechanism occurs to reduce the sensitivity of RAM meristematic cells to this hormone. A *bona fide* PM localization of MtABCG40, which functions as an importer of active forms of CK and is expressed in RAM cells (Figs 3, 4D), implies that the activity of this transporter may lower the concentration of apoplastic CKs in the root meristematic zone. As a consequence, CK free bases bind less frequently to the PM cytokinin receptors of RAM cells, the presence of which was recently confirmed by Kubiasova et al. (2020). As a result, the CK inhibitory effect on RAM could be reduced (Fig. 8). In this scenario, the role of MtABCG40 resembles that of Arabidopsis PUP14 (Zurcher et al., 2016). This statement is strengthened by the lack of changes in the total amount of active CKs and the increased expression of *MtRR4*, a primary response marker for CKs (Gonzalez-Rizzo et al., 2006), in the RAMs of *mtabcg40* compared to WT (Fig. 5C-E). Moreover, the potential increase in free base CKs in the apoplast of *mtabcg40* was also reflected by a shorter root tip in relation to WT (Fig. 5A,B) and led to shorter roots in the mutants (Fig. 2B), which was also reported for mutants in PUP14 (Zurcher et al., 2016). In *mtabcg40* roots, we observed an accumulation of IAA (Fig. 5F) and an elevation of auxin signalling (Fig. 5G). This could result from enhanced auxin biosynthesis, which also occurred after treatment with CKs (Jones et al., 2010; Yan et al., 2017), and/or a disruptive influence of CKs on polar auxin transport (Marhavy et al., 2011). Interestingly, auxin accumulation in the reduced RAMs of *mtabcg40* is consistent with the root shortening observed after exogenous IAA treatment (Fig. S1C).

**Fig. 8.**
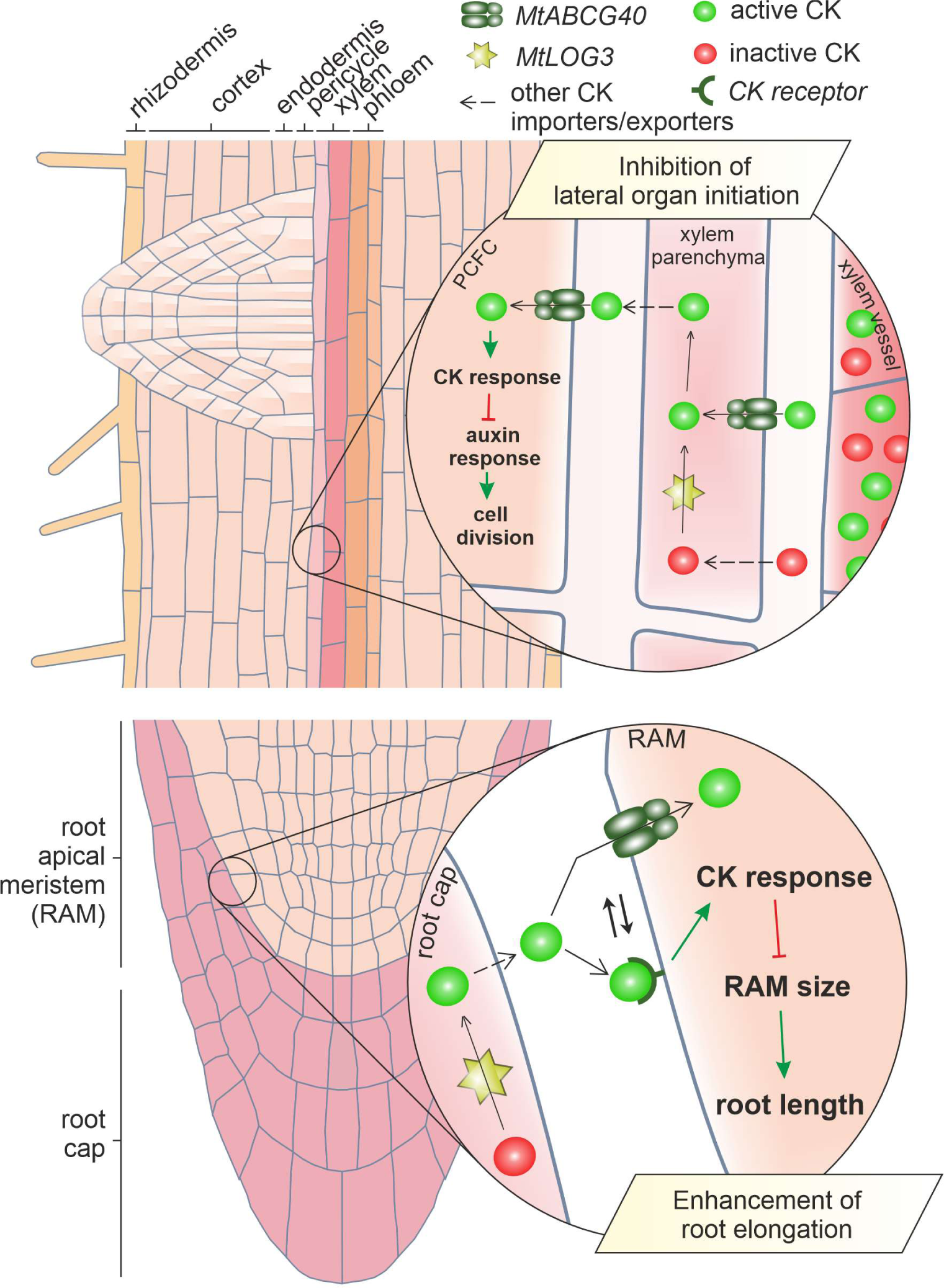
Proposed model of MtABCG40 action in *Medicago truncatula* roots. In the root vascular bundles, the root cap-biosynthesized CKs, which are biologically active and inactive, are translocated through the xylem vessels towards the shoot due to the transpiration stream. In the nitrogen-depleted environment, active CKs are imported from the apoplast surrounding the vessels to the neighbouring stele parenchyma cells by MtABCG40 (translocation mode). The latter contributes to horizontal translocation of the hormone to the cells of the pericycle, endodermis and inner cortex, which are known to generate lateral roots and nodules in *M. truncatula*, suppressing their initial divisions by inhibiting the auxin response. The translocation mode most likely requires other membrane transporters and represents a complex regulatory process considering environmental cues, such as nitrogen status. Additionally, MtABCG40-driven CK import has a more direct negative effect on auxin signalling (direct inhibitory mode) in particular cells, such as primordial contributing formation cells (PCFCs). Concomitantly, for the acquisition of the inhibitory character, inactive CKs must be first transformed by the enzyme encoded by *MtLOG3*, which is also expressed in the root vascular bundle as well as in the root cap, to their active forms. *MtLOG3* expression is induced in the root cap under nitrogen deprivation. To reduce the sensitivity of the root apical meristem (RAM) to CK, which drives root elongation in these conditions, active CKs are imported into RAM cells from the apoplast by MtABCG40. Therefore, the hormone binds to the plasma membrane CK receptors less frequently, which suppresses its characteristic outcomes, such as the negative influence on RAM size.

Apart from the regulation of primary root length, CK also influences lateral root development (Werner et al., 2001; To et al., 2004; Gonzalez-Rizzo et al., 2006). In Arabidopsis, CKs negatively impact auxin signalling in xylem pole pericycle (XPP)-derived lateral root founder cells (LRFCs), thus perturbing their transition to mitosis and suppressing the subsequent development of lateral root primordia (Laplaze et al., 2007). It was shown that in Medicago ssp. lateral roots are initiated from the pericycle, but also from endodermis and inner cortex (Xiao et al., 2019). Pericyclic cells divide prior to any other cell layers, which could point to their superior role in LR initiation (Herrbach et al., 2014). However, there is no evidence that present in the pericycle primordium contributing formation cells (PCFCs) differ from the endodermis and inner cortex cells, e.g., in their sensitivity to auxin, which would resemble the characteristics of founder cells in Arabidopsis (Dubrovsky et al., 2008; De Rybel et al., 2010). The increased number of lateral roots in *mtabcg40* (Fig. 2C) implies that MtABCG40 plays a regulatory role in the control of auxin signalling during lateral root formation. In line with this, we observed an increase in the pace of cell division (Fig. 6A,B) and a simultaneous enhancement in *MtLBD16* expression (Fig. 6D) in *mtabcg40* roots after lateral root induction using gravitropic stimulation. MtABCG40 is predominantly present inside the root stele (Figs 4D, 6C). Since xylem vessels are the source of CKs (Osugi et al., 2017), we propose that MtABCG40 imports CK free bases from the xylem apoplast to root stele cells. Moreover, prior to import by MtABCG40, the inactive CKs in the xylem should first be converted to their free bases by LOG enzymes, which are present in the root vasculature; in addition, the activity of these enzymes reduces lateral root number (Kurakawa et al., 2007) (Fig. 4). Thus, during inhibition of lateral root initiation, import of active CKs into the stele cells by MtABCG40 may facilitate an overall outwards translocation of the hormone from the vascular vessels, providing the control mechanism of CK flux toward PCFC (Fig. 8). Lateral root initiation is highlighted by the expression of *MtLBD16*, which gradually decreases during primordium development (Schiessl et al., 2019). Our observations of strong initial increase of *MtLBD16* expression in gravistimulated *mtabcg40* roots (Fig. 6D) suggest that MtABCG40 affects PCFCs initial divisions rather than LR outgrowth. In addition to the regulatory effect on overall outwards CK translocation, *MtABCG40* expression in particular stele cells, potential PCFCs (Fig. 6C), might have a more direct negative impact on auxin-stimulated cell division.

This dedicated presence of MtABCG40 as a cytokinin importer in certain cells can further adapt root patterning to external signals. In Arabidopsis, differentiated LRFCs lack PM-localized CK receptors (Kubiasova et al., 2020), thus the effect on lateral root initiation depends on intracellular CK receptors. The subcellular topology of cytokinin signalling is only partially resolved, but many cytokinin receptors are located in the ER membrane (for review see Romanov et al., 2018); therefore, ER localization can be considered in proposed here scenario.

Nodulation and lateral root formation are often considered as competing processes. This is partially due to the contrary phenotypes observed for CK-related mutants, as observed for the CK receptor *cre1* from *M. truncatula*, which exhibits decreased nodulation and increased lateral root formation (Gonzalez-Rizzo et al., 2006; Laffont et al., 2015). The difference likely results from the spatial changes in *MtCre1* expression from the pericycle and endodermis to the cortex, which occur during the transition of the root from the nonsymbiotic to symbiotic state (Boivin et al., 2016; Jardinaud et al., 2016). The latter involves the production of cortical CKs as an early prerequisite for nodule initiation (Murray et al., 2007; Reid et al., 2017). Therefore, mutating CK receptors but also transporters in the cortex inhibits nodulation (Gonzalez-Rizzo et al., 2006; Jarzyniak et al., 2021). In the nonsymbiotic state, MtABCG40 exerts a negative impact on lateral root number (Fig. 2C), similar to MtCre1 (Gonzalez-Rizzo et al., 2006). Since both genes are expressed in the vascular bundle under nonsymbiotic conditions (Boivin et al., 2016) (Fig. 4D), their corresponding phenotypes imply the presence of a mechanism that utilizes vascular CKs and suppresses lateral root development. The role of CKs in this process is further supported by an increase in lateral root number after root tip removal (Lloret et al., 1988), which eliminates a CK source in the root and the xylem vessels (Aloni et al., 2006). Interestingly, root tip removal also leads to an increase in nodule numbers (Nutman, 1952), suggesting that this vascular regulatory mechanism affects both organs in the same manner (Fig. 8). Notably, we observed an increased number of nodules, in addition to more lateral roots, in *mtabcg40* plants (Fig. 7B). Mutation in *MtABCG40* also resulted in a larger number of cell divisions (Fig. 7E,F) and an increased expression of the auxin-responsive gene *MtLBD16* (Fig. 7D) in the mutant nodule primordia. Importantly, the vascular pattern of *MtABCG40* expression did not extend to the cortex in response to symbiotic bacteria (Fig. S11). It is worth noticing that in contrast to, e.g. *MtABCG56*, expression of *MtABCG40* was not induced quickly after bacterial inoculation but at later time points (Fig. 7A). Therefore, MtABCG40 is not likely involved in early CK signalling, which promotes nodulation (Reid et al., 2017), as the MtCre1 or MtABCG40 paralogue MtABCG56 (Gonzalez-Rizzo et al., 2006; Jarzyniak et al., 2021). Notably, depending on the nitrogen status, shoot-derived and phloem-transported CKs also inhibit nodule initiation. This inhibitory effect is part of systemic autoregulation of nodulation (AON), which enables the host plant to strictly control the number of nodules (Takahara et al., 2013). Our data demonstrate that the negative regulatory role of MtABCG40 is rather local in nature and does not rely on shoot-dependent components of AON (Fig. S10). Taken together, our results demonstrate the presence of a CK-dependent inhibitory mechanism that suppresses the initiation of nodules and lateral root formation (Fig. 8). Thus, MtABCG40 integrates well into the recently suggested developmental overlaps between these two processes (Schiessl et al., 2019).

## Acknowledgements

We are thankful to J. D. Murray and E. Martinoia for critical reading; T. Ostrowski and G. Framski for the provision of chemicals; D. Weijers for the pPLV11_v02 binary vector; A. van Zeijl for *S. meliloti* 2011/pMH682; B.G. Rolfe for *S. meliloti* 1021/pHC60 strain; M.J. Barnett for *S. meliloti* A2101/pE65 strain; Sharon R. Long for *S. meliloti* Rm1021/pXLGD4 and Ora Hazak for the binary *HDEL-mCherry* plasmid. This work was supported by the National Science Centre, Poland (grant no. 2015/19/B/NZ9/03548 to M.J.), the Swiss National Funds (project 310030_197563 to M.G.), and the ERDF project “Plants as a tool for sustainable global development” (No. CZ.02.1.01/0.0/0.0/16_019/0000827 to O.N.).

## Author contributions

M.J. devised and supervised the project. T.J., M.J. and J.B. designed the experiments and interpreted the results. T.J. performed the majority of the experiments. T.J. and A.P. performed phenotypic characterization of plants, including mutants and silenced material. J.B. and T.J. performed microscopic observations (RAM length, EdU staining, promoter analyses). T.J., A.P. and K.J. generated the plasmids and performed qRT-PCR analyses. T.J. and A.P. performed plant transformation. M.G. designed and T.T., F.R.I. and J.X performed transport experiments. F.R.I performed subcellular localization in *N. benthamiana* leaf epidermal cells. O.N., W.B.L. and L.P. conducted quantification of endogenous cytokinins and auxins. T.J. and J.B. conducted statistical analyses. T.J. and J.B. prepared figures. T.J. proposed a working model. T.J., M.J. and J.B. wrote the manuscript with the help of co-authors. All authors saw and commented on the manuscript.

## Data availability

All data are included in the main article or the Supporting Information. The sequence data from this article can be found in Phytozome v13 database under the following accession numbers: *MtABCG40* (Medtr7g098300), *MtLOG1* (Medtr7g101290), *MtLOG2* (Medtr1g064260), *MtLOG3* (Medtr1g057020), *MtLOG-like 1* (Medtr1g015830), *MtLOG-like 2* (Medtr2g094790), *MtLOG-like 3* (Medtr3g113710), *MtLOG-like 4* (Medtr4g058740), *MtLBD16* (Medtr7g096530), *MtRR4* (Medtr5g036480), *Mtactin* (Medtr3g095530), *MtTML2* (Medtr6g023805).

## SUPPORTING INFORMATION

Fig. S1 Primary root length and lateral root number of *Medicago truncatula* at different concentrations of ammonium nitrate (NH_4_NO_3_), indole-3-acetic acid (IAA), and 6- benzylaminopurine (6-BAP).

Fig. S2 Characterization of NF21323 (*mtabcg40-1*) and NF17891 (*mtabcg40-2*) mutant lines of *MtABCG40* used in the study.

Fig. S3 Lateral root density of WT and mutant (*mtabcg40*) plants grown on media supplemented with 1 mM NH_4_NO_3_.

Fig. S4 Phylogenetic tree of full-size ABCG proteins from *Arabidopsis thaliana* and *Medicago truncatula* showing the close relation of MtABCG40 and MtABCG56.

Fig. S5 Co-expression of MtABCG40 with an ER marker and transport controls.

Fig. S6 A heat map showing a decline in *AtLOG7* expression in the root pericycle triggered within 48 h after an addition of 5 mM NH_4_NO_3_ to the nitrogen-depleted (0.3 mM NH_4_NO_3_) media.

Fig. S7 Comparison of the size of RAM between WT-1 and *mtabcg40-1* mutant plants grown under nitrogen-sufficient conditions (medium supplemented with 1 mM NH_4_NO_3_).

Fig. S8 Schematic representation of lateral root induction using gravitropic stimulation.

Fig. S9 Comparison of the cell division rate and *MtLBD16* expression within gravitropically- stimulated lateral root primordium between WT-1 and *mtabcg40-1* mutant plants grown under nitrogen-sufficient conditions (medium supplemented with 1 mM NH_4_NO_3_).

Fig. S10 Analysis of a systemic control of *MtABCG40* expression during autoregulation of nodulation (AON) using *Medicago truncatula* plants with split-root system.

Fig. S11 Expression pattern of MtABCG40 during different stages of the symbiotic interaction.

Table S1 List of primers used in the study.

**Excel File S1** Details regarding statistical analyses.

