## supplementary file for "*Medicago truncatula* ABCG40 is a cytokinin importer that negatively regulates lateral root density and nodule number"

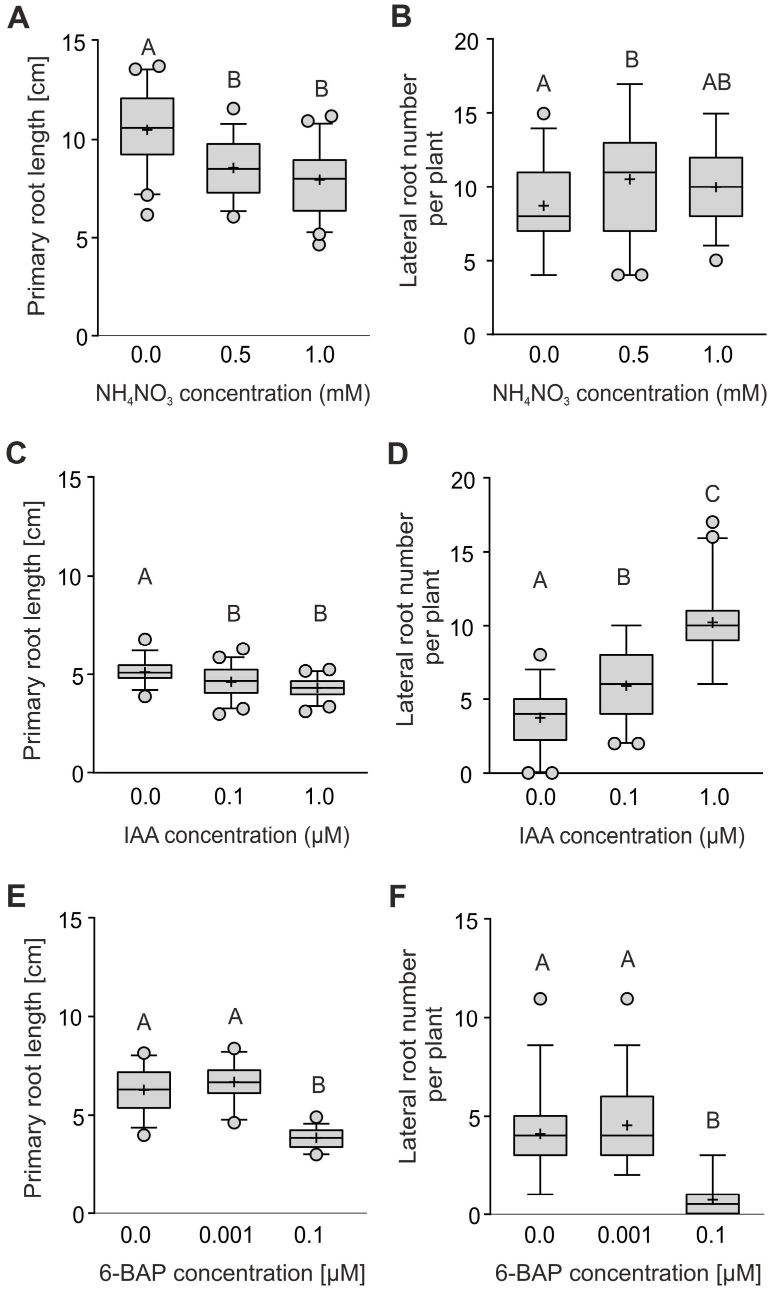


**Fig. S1** Primary root length and lateral root number of *Medicago truncatula* grown under different concentrations of ammonium nitrate ($NH_4NO_3$) (A and B), indole-3-acetic acid (IAA) (C and D), and 6-benzylaminopurine (6-BAP) (E and F). The box plots present the main root length and lateral root number for 31-45 roots per condition obtained from three independent biological experiments. For each box-and-whiskers plot: the central black line represents the median; '+' represents the mean; the box extends from the 25th to 75th percentiles; the whiskers are drawn down to the 5th percentile and up to the 95th. Points below and above the whiskers are drawn as individual dots. Identical or different uppercase letters indicate no or significant differences, respectively; $P < 0.05$. Significant differences were determined by one-way ANOVA with a post hoc Tukey's multiple comparison test (A), Kruskal–Wallis test with a post hoc Dunn's multiple comparison test (B, D and F), and Brown-Forsythe and Welch ANOVA with a post hoc Tamhane's T2 multiple comparison test (C and E).

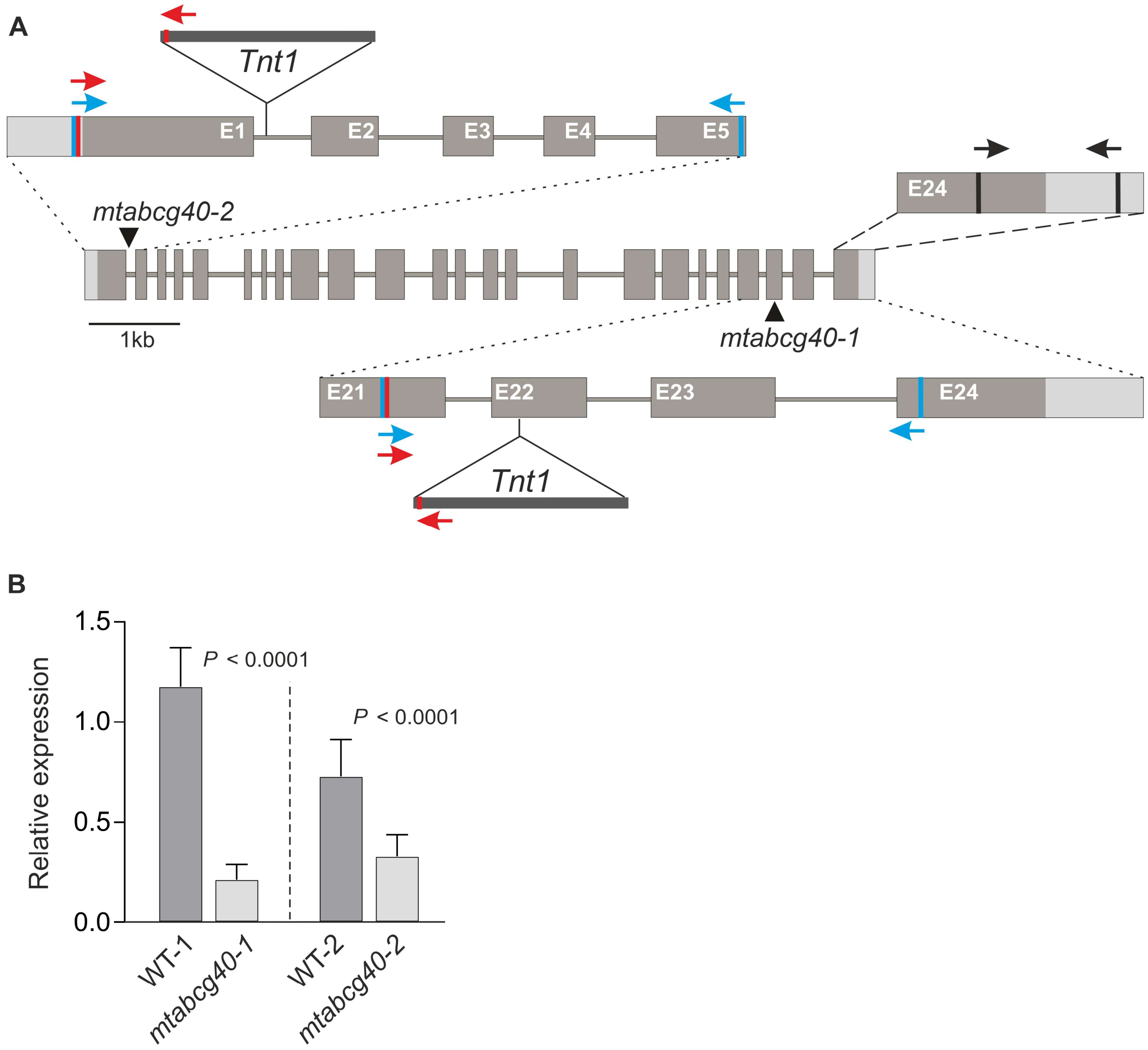


**Fig. S2** Characterization of NF21323 (*mtabcg40-1*) and NF17891 (*mtabcg40-2*) mutant lines of *MtABCG40* used in the study. A, Schematic representation of *MtABCG40* gene structure with localization of *Tnt1* insertions indicated by triangles. Grey boxes and the gaps between the boxes indicate exons and introns, respectively. Blue and red arrows indicate location of primers used for genotyping of WT and mutant alleles, respectively. Black arrows indicate location of primers used for the determination of *MtABCG40* transcript accumulation in WT and mutant lines. B, *MtABCG40* transcript accumulation in the roots of control (WT) and mutant (*mtabcg40*) plants upon 0 mM $NH_4NO_3$. Transcript levels were measured by quantitative real-time PCR and normalized to the *Mtactin*. Expression data represent the mean ± SD of three independent biological experiments and two or three technical repeats. Significant differences from the WT plants were determined by two-tailed Student's t-test with Welch correction.

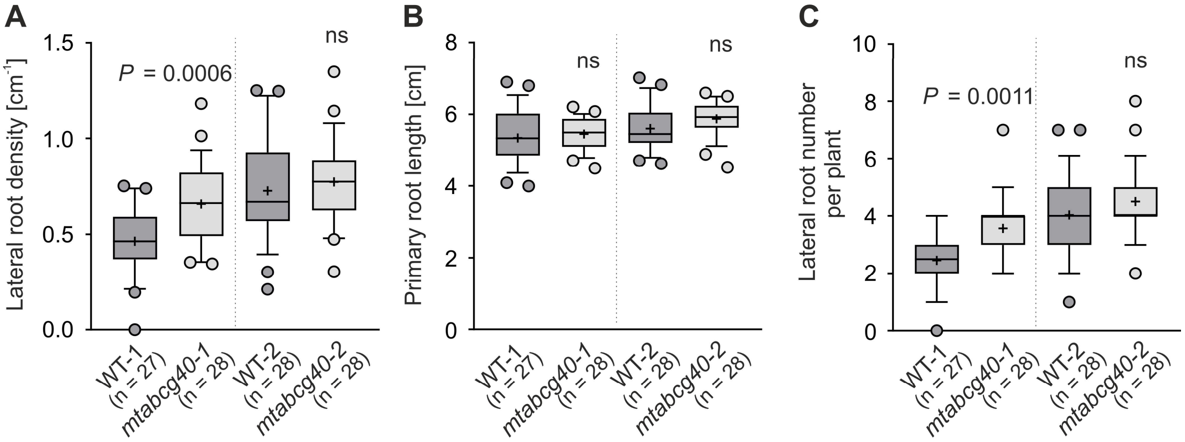


**Fig. S3** Lateral root density of WT and mutant (*mtabcg40)* plants grown on media supplemented with 1 mM $NH_4NO_3$. For each box-and-whiskers plot: the central black line represents the median; '+' represents the mean; the box extends from the 25th to 75th percentiles; the whiskers are drawn down to the 10th percentile and up to the 90th. Points below and above the whiskers are drawn as individual dots. Significant differences from the WT control were determined by two-tailed Student's t-test (A), two-tailed Student's t-test with Welch correction (B), two-tailed Mann–Whitney test (C); ns, not significant; n represents the number of individual roots obtained from two independent biological experiments.

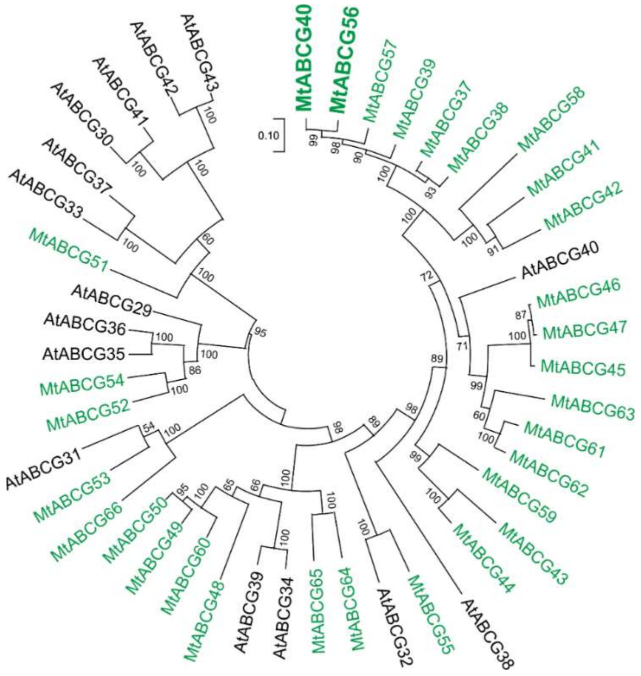


**Fig. S4** Phylogenetic tree of full-size ABCG proteins from *Arabidopsis thaliana* (Verrier et al., 2008) and *Medicago truncatula* (Jarzyniak et al., 2021) showing the close relation of MtABCG40 and MtABCG56. The evolutionary history was inferred using the Maximum Likelihood method and JTT matrix-based model (bootstraps: 1000) (Jones et al., 1992) based on the amino acid sequences generated after multiple sequence alignments with MUSCLE. The tree with the highest log likelihood (-46855.58) is shown. The tree is drawn to scale, with branch lengths measured in the number of substitutions per site. This analysis involved 45 amino acid sequences. All positions containing gaps and missing data were eliminated (complete deletion option). There were a total of 1194 positions in the final dataset. Evolutionary analyses were conducted in MEGA X (Kumar et al., 2018)

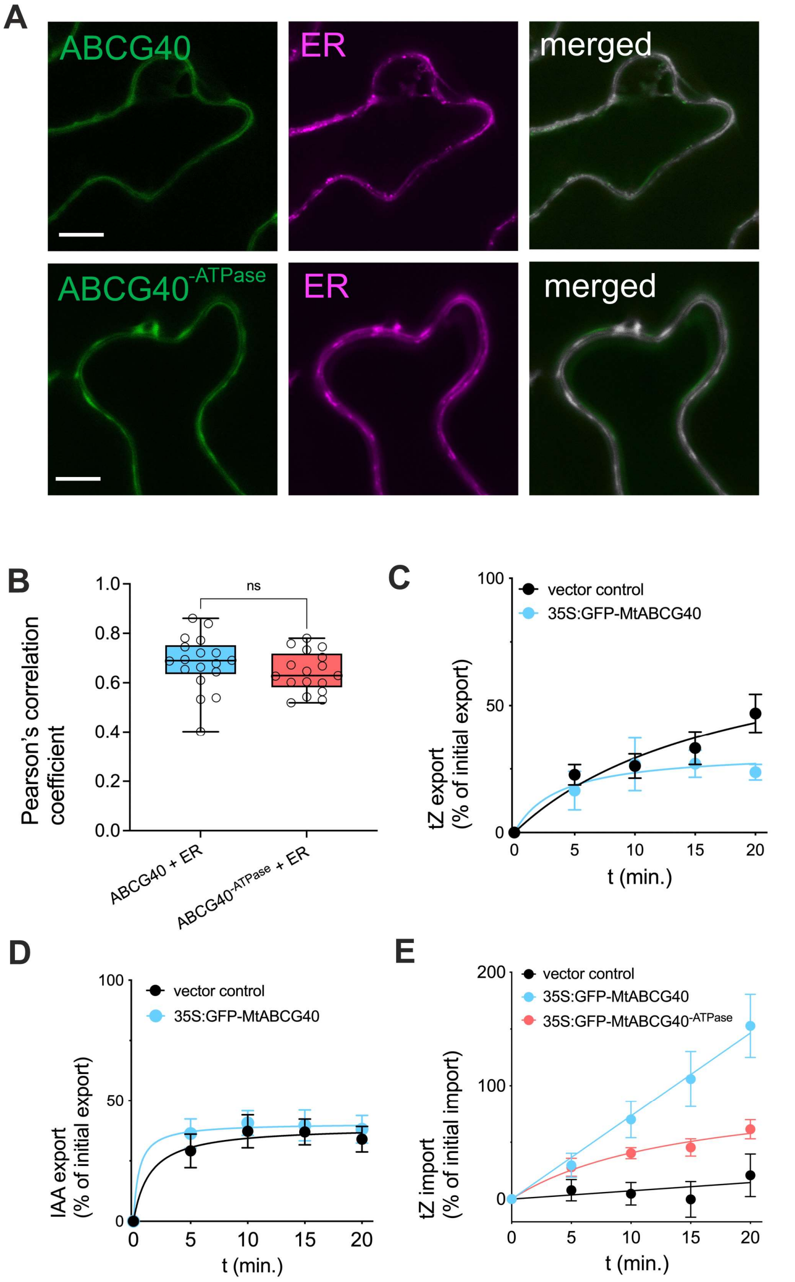


**Fig. S5** Co-expression of MtABCG40 with an ER marker and transport controls. A and B, Co-expression of GFP-MtABCG40 and GFP-MtABCG40$^{-ATPase}$ fusion proteins in Agrobacterium-infiltrated *Nicotiana benthamiana* leaf epidermal cells with the ER marker, HDEL-mCherry. Confocal imaging (A) and Pearson's correlation coefficients (B). GFP-MtABCG40, GFP-MtABCG40$^{-ATPase}$ and ABCB1-RFP images were pseudo-colored in green and magenta, respectively. Scale bars, 50 µm. C-E, Transport experiments in tobacco protoplasts expressing *GFP-MtABCG40* and *GFP-MtABCG40$^{-ATPase}$*. Relative [$^{14}$C]-*t*Z (C) and [$^{3}$H]-indole-3-acetic acid (IAA) export (D) as well as [$^{14}$C]-*t*Z import (E) from tobacco protoplasts, as well as into tobacco protoplasts. Data represent the result from a minimum of 4 independent experiments (transfections and protoplast preparations).

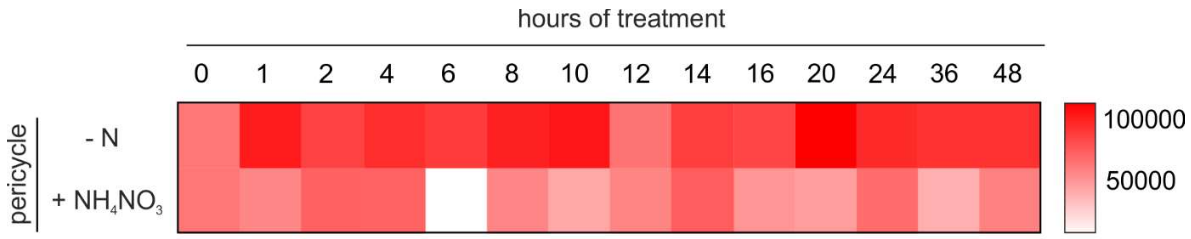


**Fig. S6** A heat map showing a decline in AtLOG7 expression in the root pericycle triggered within 48 h after an addition of 5 mM NH4NO3 to the nitrogen-depleted (0.3 mM NH4NO3) media. The graph depicts log10-fold changes. The transcriptomic data acquired from (Walker et al., 2017).

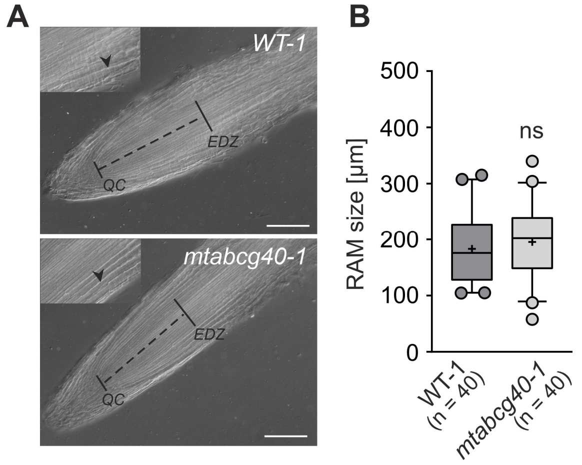


**Fig. S7** Comparison of the size of root apical meristem (RAM) between WT-1 and *mtabcg40-1* mutant plants grown under nitrogen-sufficient conditions (medium supplemented with 1 mM $NH_4NO_3$). A, Nomarski image of the RAM and apical region of the elongation/differentiation zone (EDZ) of a primary root 10 days after germination. The size of the RAM (black dotted line) was determined as the distance from the quiescent center (QC) to the EDZ. The apical border of the EDZ was defined by the first elongated cortical cell of the second cortical layer (arrowhead). Scale bars, 100 µm. B, Graph showing the RAM size of *mtabcg40-1* mutant and WT-1. For each box-and-whiskers plot: the central black line represents the median; '+' represents the mean; the box extends from the 25th to 75th percentiles; the whiskers are drawn down to the 5th percentile and up to the 95th. Points below and above the whiskers are drawn as individual dots; n represents the number of individual plants obtained from two independent experiments. Significant differences from the control plants (WT-1) determined by two-tailed Mann–Whitney test; ns, not significant.

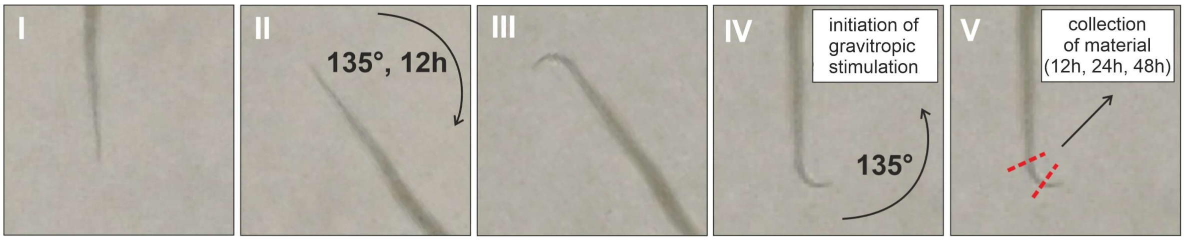


**Fig. S8** Schematic representation of lateral root induction using gravitropic stimulation. 3-day-old seedlings (step I) are turned 135° for 12 h (step II) to induce the growth-mediated curvature of the root (step III) and turned back afterwards to the initial orientation to induce lateral root morphogenesis through gravitropism (step IV). Samples comprising bent fragments are collected in an experiment-specific time points after an induction for further analysis (step V).

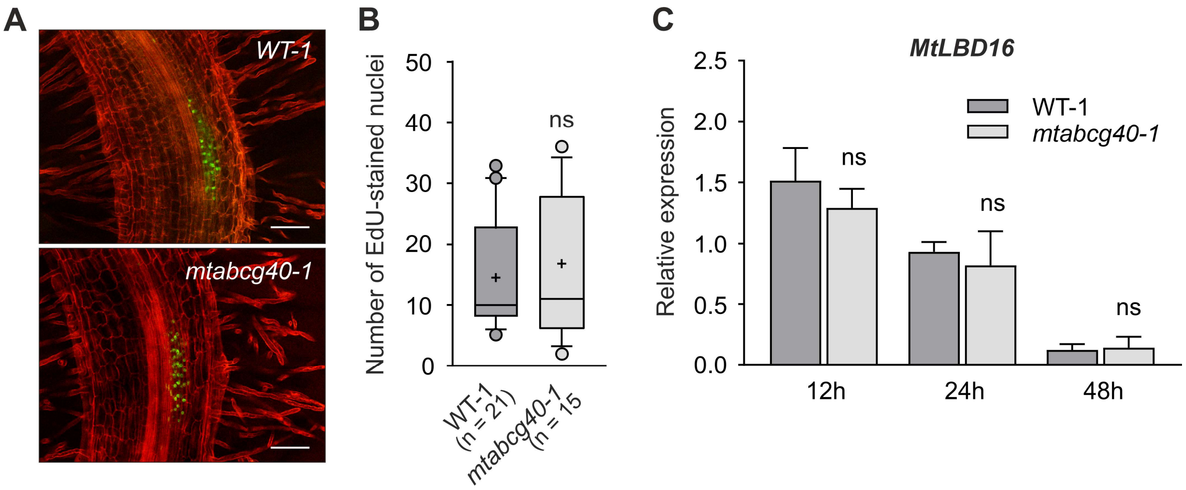


**Fig. S9** Comparison of the cell division rate and *MtLBD16* expression within gravitropically-stimulated lateral root primordium between WT-1 and *mtabcg40-1* mutant plants grown under nitrogen-sufficient conditions (medium supplemented with 1 mM $NH_4NO_3$). A, Optical sections of lateral root primordium 12 hours post induction (hpi) by gravitropic stimulation. Red propidium iodide demarks cell walls and green EdU-labeled nuclei DNA replication. Scale bars, 100 µm. B, Comparison of the number of EdU-stained nuclei within lateral root primordium between WT-1 and the *mtabcg40-1* mutant. For each box-and-whiskers plot: the central black line represents the median; '+' represents the mean; the box extends from the 25th to 75th percentiles; the whiskers are drawn down to the 5th percentile and up to the 95th. Points below and above the whiskers are drawn as individual dots; n represents the number of individual plants obtained from two independent experiments. C, Expression analysis of the *MtLBD16* within the gravitropically stimulated bent fragments (2-3 mm) at indicated time points, 12, 24 and 48 hpi. Transcript levels measured by quantitative real-time PCR and normalized to *Mtactin*; data represent the mean ± SD of two independent biological experiments with three technical repeats each. Significant differences from the control plants (WT-1) determined by two-tailed Mann–Whitney test (B and C); ns, not significant.

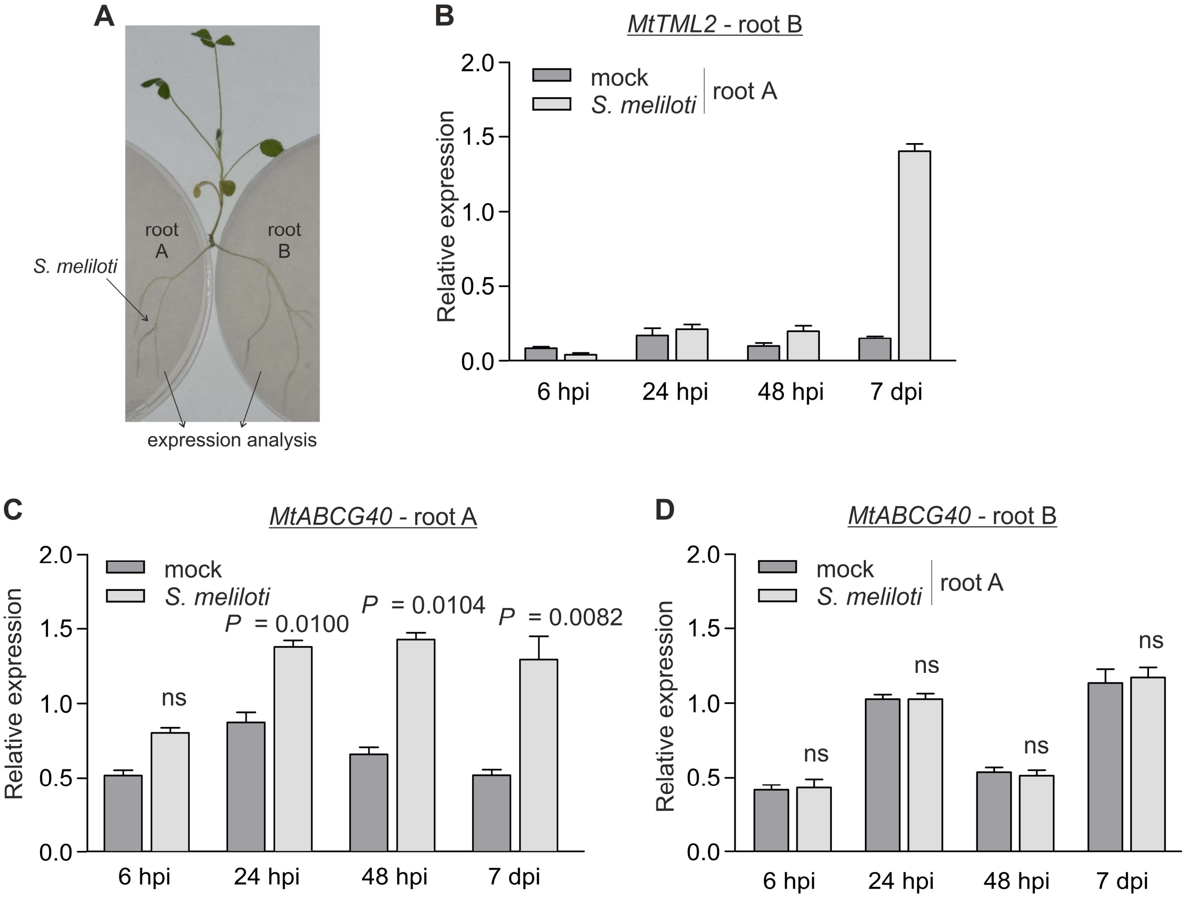


**Fig. S10** Analysis of a systemic control of *MtABCG40* expression during autoregulation of nodulation (AON) using *Medicago truncatula* plants with split-root system. A, Schematic representation of an AON analysis. Plants were grown for 3 weeks on N-depleted medium with both roots spatially separated. Subsequently, one of these roots, named A, was flood inoculated with *Sinorhizobium meliloti* suspension. Both roots, A and not inoculated B, were then collected at specified time points and subjected to gene expression analysis. Roots 6, 24, 48 hours post inoculation (hpi) and 7 days post inoculation (dpi) were analyzed. B, Expression analysis of *MtTML2*, a AON marker gene in root B. C and D, Expression analysis of *MtABCG40* in root A (C) and B (D). Transcript levels were measured by quantitative real-time PCR and normalized to the *Mtactin*. Expression data represent the mean ± SD of three technical repeats. Significant differences from the control plants determined by two-tailed Mann–Whitney test.

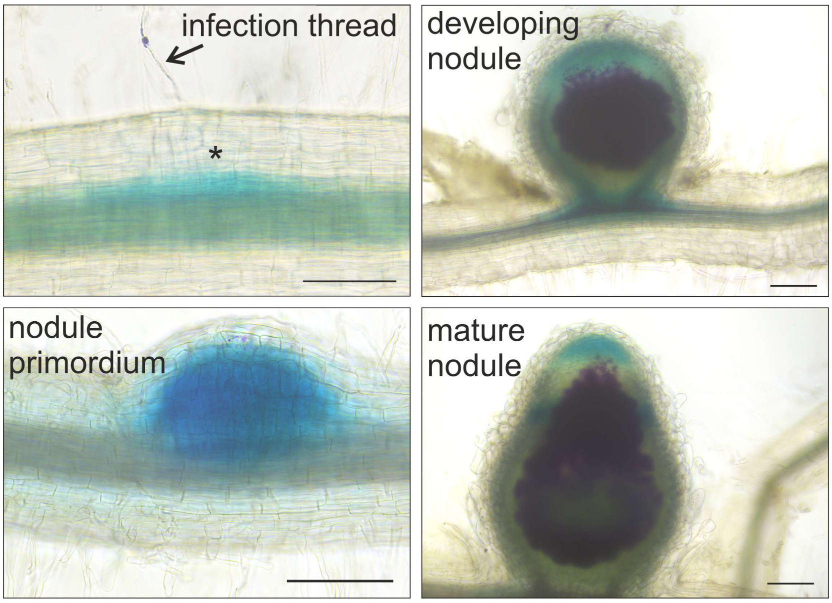


**Fig. S11** Expression pattern of *MtABCG40* during different stages of the symbiotic interaction. Composite transgenic plants carrying *ProMtABCG40:GUS* construct were imaged at different stages after inoculation with *Sinorhizobium meliloti* Rm1021/pXLGD4 (lacZ). Double staining using Magenta-Gal and X-Gluc allowed the visualization of the infecting *S. meliloti* in magenta and *MtABCG40* expression in blue. Scale bars, 100 µm.

### **Table S1** List of primers used in the study

A. Primers used for the identification of *Tnt1* mutant lines

| **Primer name** | **Forward primer (5' → 3')** | **Reverse primer (3' → 5')** |
|---|---|---|
| *mtabcg40-1* M | AATGTATGGCTTGCTTATGG | GCTACCAACCAAACCAAGTC |
| MtABCG40-1 WT | AATGTATGGCTTGCTTATGG | ATCTCCATATTGTGAAGCCG |
| *mtabcg40-2* M | TAACTGGGGTATCGTAAAGG | GCTACCAACCAAACCAAGTC |
| MtABCG40-2 WT | TAACTGGGGTATCGTAAAGG | CATATTGAGGTCCAACTCCTTGG |
| *MtABCG40* qPCR | CAATGATGGAAGCCAAACC | CTTTATAGAAGCCGTATCAC |

B. Primers used for the analyses of promoter activity (gene specific sequences underlined)

| **Primer name** | **Forward primer (5' → 3')** | **Reverse primer (3' → 5')** |
|---|---|---|
| *ProMtABCG40: GUS* | ggggacaagtttgtacaaaaaagcaggcTTCAGTAACGAATATGGTAGG | ggggaccactttgtacaagaaagctgggtCAATAACAGGAAAAAGTTTTATCC |
| *ProMtABCG40: NLS-GFP* | tagttggaatgggttcgaaTCAGTAACGAATATGGTAGG | ttatggagttgggttcgaaCAATAACAGGAAAAAGTTTTATCC |
| *ProMtLOG3: GUS* | ggggacaagtttgtacaaaaaagcaggcTCTAAAATTGCTAAGTTCAGAGG | ggggaccactttgtacaagaaagctgggtGATCTTCTTTGTGGAAAGGT |
| *ProMtLOG3: NLS-tdTomato* | tagttggaatgggttcgaaAATGATTTTATAGATGTGTCGG | ttatggagttgggttcgaaGATCTTCTTTGTGGAAAGG |

C. Primers used for RNAi silencing (gene specific sequences underlined)

| **Primer name** | **Forward primer (5' → 3')** | **Reverse primer (3' → 5')** |
|---|---|---|
| *MtABCG40* RNAi | ggggacaagtttgtacaaaaaagcaggctAATATGTAGCATAAACATATGAACA | ggggaccactttgtacaagaaagctgggtCAATAACAGGAAAAAGTTTTATCC |

| | | |
|---|---|---|
| *MtLOG3* RNAi | ggggacaagtttgtacaaaaaagcaggctTGATC TTATATAAGTAAGACTAGTGG | ggggaccactttgtacaagaaagctgggtGCATCATCA ACATTAACATTAGTTG |

D. Primers for Real-Time PCR analyses

| **Primer name** | **Forward primer (5’ → 3’)** | **Reverse primer (3’ → 5’)** |
|---|---|---|
| *MtABCG40* | CAA TGA TGG AAG CCA AAC C | CTTTATAGAAGCCGTATCAC |
| *MtLOG1* | ACTAACAAATGGACTCACGC | CTCCAGAGTTCCATATCCC |
| *MtLOG2* | GAGAGCTAACTGGTGAAACG | GCATTTGGACTGATAAATCCC |
| *MtLOG3* | AAGAGAGATAACTGGTGACC | CTCTAATTCCCTAACCAATTCC |
| *MtLOG3* RNAi | TGATCTTATATAAGTAAGACTAGTGG | GCATCATCAACATTAACATTAGTTG |
| *MtLOG-like 1* | GAACTATGGAAGAGCTTCTGG | GGAAGGAGAGTAACTCTCCA |
| *MtLOG-like 2* | ATTGCTGAATGTTGATGGATAC | CTCTACACAAACTGCCATCG |
| *MtLOG-like 3* | ATGAAGGTTTTGTAACACCAGC | GTTCAAAGTAGAAAATCAGCGG |
| *MtLOG-like 4* | AGAGATCACTGGAGAGACAG | AGCCATCTACATTCAACAACC |
| *MtRR4* | CTGAATCTGATGCTTTTGTTCC | CCTCCAAACATCTGTCAATGC |
| *MtLBD16* | AGCTCGTATCAGAGACCCTGTG | TGCAAGCATGCTACCTGTTGTTG |
| *Mtactin* | AAGCATCACAATCACTCC | TTCTCTCAGTACTTTCCAGC |
| *MtTML2* | TCTGGTGACAATGGTTCCTC | AAGACATGGTAATGGTAGTAGAAC |
